# Cephalosporins target quorum sensing and suppress virulence of *Pseudomonas aeruginosa* in *Caenorhabditis elegans* infection model

**DOI:** 10.1101/2020.05.15.097790

**Authors:** Lokender Kumar, Nathanael Brenner, John Brice, Judith Klein-Seetharaman, Susanta K. Sarkar

## Abstract

*Pseudomonas aeruginosa* utilizes a chemical social networking system referred to as quorum sensing (QS) to strategically co-ordinate the expression of virulence factors and biofilm formation. Virulence attributes damage the host cells, impair the host immune system, and protect bacterial cells from antibiotic attack. Thus, anti-QS agents may act as novel anti-infective therapeutics to treat *P. aeruginosa* infections. The present study was performed to evaluate the anti-QS, anti-biofilm, and anti-virulence activity of β-lactam antibiotics (carbapenems and cephalosporins) against *P. aeruginosa*. The anti-QS activity was quantified using *Chromobacterium violaceum* CV026 as a QS reporter strain. Our results showed that cephalosporins including cefepime (CP), ceftazidime (CF), and ceftriaxone (CT) exhibited potent anti-QS and anti-virulence activities against *P. aeruginosa* PAO1. These antibiotics significantly impaired motility phenotypes, decreased pyocyanin production, and reduced the biofilm formation by *P. aeruginosa* PAO1. In the present study, we studied isogenic QS mutants of PAO1: ΔLasR, ΔRhlR, ΔPqsA, and ΔPqsR and found that the levels of virulence factors of antibiotic-treated PAO1 were comparable to QS mutant strains. Molecular docking predicted high binding affinities of cephalosporins for the ligand-binding pocket of QS receptors (CviR, LasR, and PqsR). In addition, our results showed that the anti-microbial activity of aminoglycosides increased in the presence of sub-inhibitory concentrations (sub-MICs) of CP against *P. aeruginosa* PAO1. Further, utilizing *Caenorhabditis elegans* as an animal model for the *in vivo* anti-virulence effects of antibiotics, cephalosporins showed a significant increase in *C. elegans* survival by suppressing virulence factor production in *P. aeruginosa*. Thus, our results indicate that cephalosporins might provide a viable anti-virulence therapy in the treatment of infections caused by multi-drug resistant *P. aeruginosa*.

## INTRODUCTION

*Pseudomonas aeruginosa* causes nosocomial infections and multi-drug-resistant (MDR) *P. aeruginosa* is increasingly problematic. Bacterial communication is critical for infection, virulence, survival, antibiotic resistance, and biofilm formation in *P. aeruginosa* [1–5]. Cells communicate by expressing specific chemical signal molecules that lead to coordinate their population-wide behavior, a process known as quorum sensing (QS) [4]. QS involves the expression of acyl-homoserine lactone (AHL) molecules and detecting these signal molecules using specific protein receptors [6]. In the *P. aeruginosa* QS system, homo-serine lactone autoinducer molecules (HSL) produced by LuxI-type enzymes bind to cognate transcriptional regulator LuxR-type protein receptors enabling receptor dimerization and binding to DNA promoter sequences [7]. Further, this signal–receptor complex acts as a transcriptional regulator and initiates the expression of an array of virulence and biofilm formation associated genes [8].

The *P. aeruginosa* QS network is comprised of three co-dependent systems, the las [9], rhl [10], and pqs [11] systems. Las and rhl respond to N-(3-oxododecanoyl) homoserine lactone (3O-C12–HSL) [12] and N-butyryl homoserine lactone (C4–HSL) [13]. The LasR/3OC12-HSL complex also activates transcription of rhlR system, and Rhl/C4-HSL enhances the expression of numerous genes including the las regulon genes [14]. Pqs activation results in the synthesis of 2-heptyl-3-hydroxy-4-quinolone (PQS) that plays a connecting role between the las and rhl QS systems [2]. This global regulatory organization of QS allows *P. aeruginosa* to synchronize population-wide behaviors and survive under hostile environmental conditions [15]. Emerging research has suggested that QS mutant strains form thinner biofilms and show decreased expression of virulence factors than wild type strains [16]. QS is important in regulating growth, biofilm formation and exopolysaccharide production [12, 17, 18]. Additionally, the LasR signaling molecule, 3O-C12-HSL showed immunomodulatory activities influencing the progression of *P. aeruginosa* infection [19, 20]. Finally, QS pathways may help *P. aeruginosa* to form thick biofilms protected by extracellular polysaccharide matrix (alginate) that defends planktonic cells against antimicrobial attacks [21–23].

QS inhibition has been considered as a new antimicrobial approach to attenuate the virulence of *P. aeruginosa* [24]. It can be accomplished by either signal molecule degradation or targeting interaction of the signal molecules with their cognate receptors [25]. Since this strategy does not directly target vital pathways required for bacterial survival, the probability of emergence of resistant bacterial strains against anti-QS drugs is low. Thus, targeting the QS cell-signaling pathways can offer an innovative anti-virulence strategy against *P. aeruginosa* with lower emergence of resistance. Antimicrobial agents [26] and phytochemicals [27] have shown potent anti-virulence activity by inhibiting QS pathways in *P. aeruginosa. In-vitro* phenotypic characterization and *in silico* molecular docking analysis have shown that the antifungal drugs (clotrimazole and miconazole) and antibacterial agents (for example, clofoctol) have potent anti-QS activity [28]. Phytochemicals (zingerone, gingerol, and curcumin) have strong anti-QS activity and effectively inhibit virulence and biofilm formation of *P. aeruginosa* [8, 29, 30]. These molecules have also shown the synergistic effect in combination with other antimicrobial agents to eradicate *P. aeruginosa* biofilms [25]. Furthermore, natural anti-QS molecules include halofuranes, originated from macroalgae, specifically furanone C30 strongly inhibits QS at relatively low doses (1-10 μM) [31]. Most of these compounds have shown anti-QS properties at lower concentrations and anti-microbial properties at higher concentrations [25].

Sub-inhibitory concentrations of antibiotics are believed to interfere with gene expression during the growth cycle of bacteria [32], raising the question if the antibiotics might also have alternate inhibitory pathways against pathogen bacteria. Macrolide antibiotics (azithromycin) have improved the clinical outcome of *P. aeruginosa* infection with chronic pulmonary disorders and cystic fibrosis [33, 34]. In addition, tobramycin [35], ciprofloxacin [36] and doxycycline [37] have been reported to inhibit quorum-sensing pathways in *P. aeruginosa*. The broad-spectrum cephalosporin antibiotics provide protective efficacy and improved the outcome of *P. aeruginosa* infections [38, 39]. Cefepime (CP) [40, 41], ceftazidime (CF) [42] and ceftriaxone (CT) [43] treatments have shown remarkable activity against *P. aeruginosa* infection, however, the emergence of drug resistance has restricted the use of these antibiotics against many Gram-negative pathogens including *P. aeruginosa* [44, 45]. CF, a third-generation cephalosporin antibiotic inhibits QS pathways by binding with QS receptors (LasR and RhlR) [46]. Consequently, the use of anti-QS or antivirulence antibiotics at sub-inhibitory concentrations may improve the outcome of infection and reduce the emergence of antibiotic resistance phenotypes. Based on the above findings, we propose the hypothesis that the anti-QS cephalosporin antibiotics will display potent anti-virulence potential against *P. aeruginosa* infections.

It has been shown that the model organism *Chromobacterium violaceum* utilizes AHL dependent QS pathways to produces violacein pigment to make purple bacterial colonies. The CV026 mutant (an AHL synthase deletion) appears as colorless since it cannot produce its signaling molecule. However, it can respond to AHL supplements and produce purple color colonies. The CV026 strain has been exploited to screen anti-QS compounds and the selected compounds are potential anti-QS drugs against *Pseudomonas aeruginosa*. By screening several major cephalosporins, we find that CP, CF, and CT show potent anti-QS activity against *Chromobacterium violaceum* CV026. Further, sub-inhibitory concentrations of these antibiotics displayed potent *in vitro* anti-virulence activity and synergistically enhance the antimicrobial activity of aminoglycosides against *P. aeruginosa* PAO1. Furthermore, molecular docking shows significant interactions of *C. violaceum* and *P. aeruginosa* QS receptors with cephalosporins. To corroborate these findings in an *in vivo* setting, we tested antivirulence efficacy of CP, CF, and CT in the *C.elegans* infection model, validating their anti-virulence potential against *P. aeruginosa*.

## RESULTS

### Anti-QS activity of cephalosporins against *C. violaceum* CV026

Results of agar well diffusion assay showed that cephalosporins displayed growth inhibitory zones (clear circular area surrounding wells) reporting their antimicrobial activity against *C. violaceum* CV026 **(Fig 1A-1C, Supplementary Fig 1).** In addition to the antimicrobial zone, there was a distinct halo zone of pigment inhibition at the interface of bacterial growth (purple color) and anti-microbial zone (largest among CP, CF, and CT-**Supplementary Fig 1**). The halo zone of pigment inhibition was designated as the anti-QS zone. Each antibiotic was tested in eight dilutions (4 wells in each plate) with numbers 1-8 indicating highest to lowest concentrations of antibiotics (from 512 μg to 0.4 μg). Most of the antibiotics showed a significant zone of anti-QS activity **(Supplementary Fig 2)**. The largest anti-QS zone was exhibited by CP (0.66 + 0.87 mm) **(Fig 1A)**, CF (0.48 + 0.52 mm) **(Fig 1B)**, and CT (0.24 + 0.36 mm) **(Fig 1C)**. Other cephalosporin antibiotics showed a small yet significant zone (p<0.05) of anti-QS activity including imipenem (0.19 + 0.07 mm), doripenem (0.26 + 0.87 mm), meropenem (0.25 + 0.13 mm) and ertapenem (0.31 + 0.09 mm) **(Supplementary Fig 3)**.

**Figure-1:**
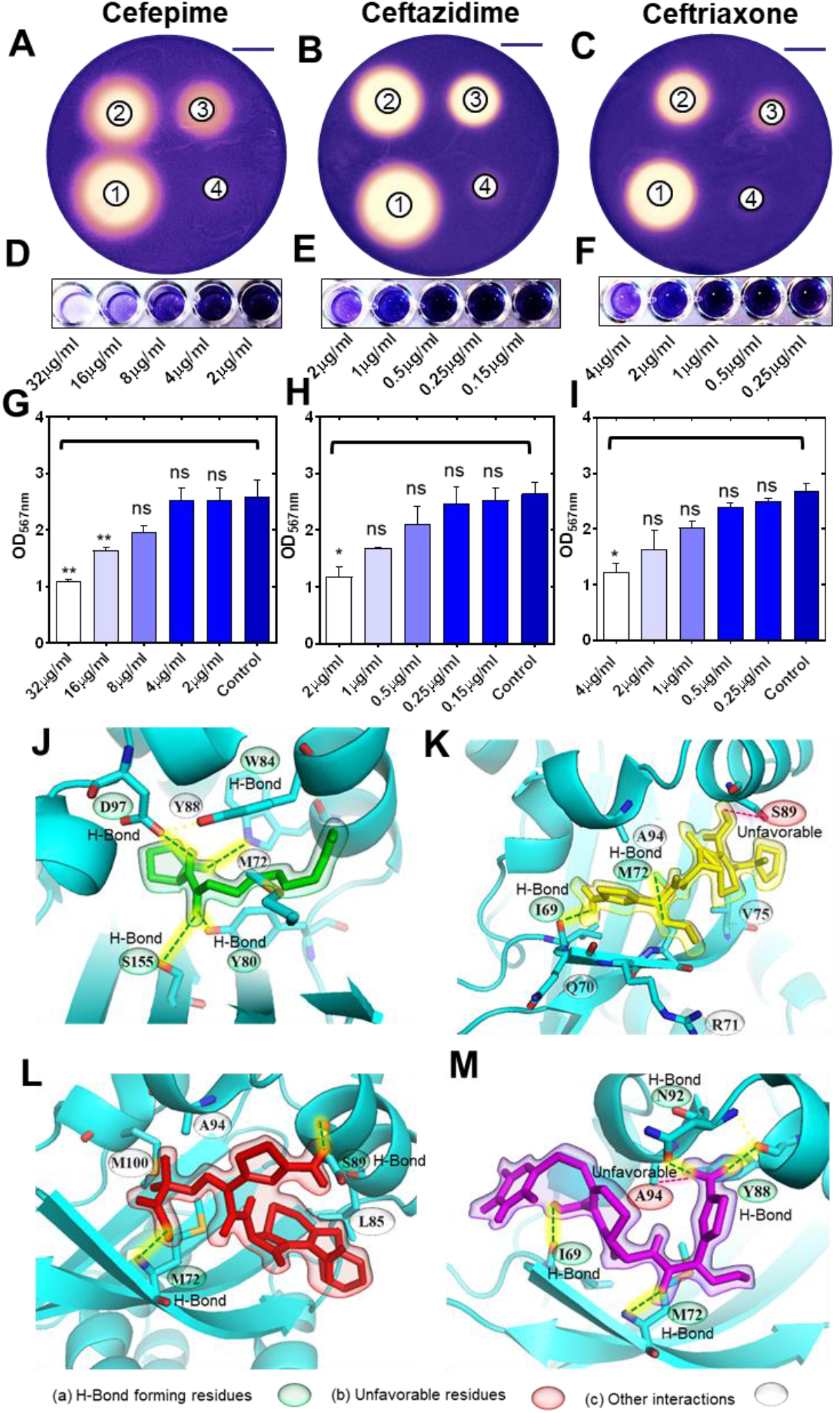
Inhibition of QS in *Chromobacterium violaceum* CV026 by cephalosporin antibiotics. Photographs of *C. violaceum* plates showing the zone of growth inhibition and pigment production inhibition (on the edge of the zones) by CP (A), CF (B) and CT (C) against *C. violaceum* CV026 (well-1= 51.2 μg; well-2=25.6 μg; well-3= 12.8 μg; well-4=6.4 μg) (Scale bar= 20mm). Images of microtiter plate wells and OD 567nm measurements showing the inhibition of pigment production in 96 well plate assay by sub-inhibitory concentrations of CP (D, G), CF (E, H) and CT (F, I). Molecular docking results showing the binding interaction of natural ligand C-10 homoserine lactone molecule (J), CP (K), CF (L), and CT (M) with CviR QS receptor of *C. violaceum* (ns p > 0.05, * p ≤ 0.05, ** p ≤ 0.01).

Further, to validate that the sub-inhibitory concentrations of CP, CF, and CT **(Supplementary Fig 4)** were responsible for pigment inhibition, we determined the minimum inhibitory concentration (MIC) of these cephalosporin antibiotics **(Supplementary Fig 5A-5D)** against the CV026 strain. MICs of CP, CF, and CT against CV026 were 128 μg/mL, 8 μg/mL and 16 μg/mL **(Supplementary Fig 5A**). It was observed that the growth of CV026 was significantly inhibited at sub-MIC concentrations of antibiotics indicating substantial effects of antibiotics on molecular pathways of bacterial growth **(Supplementary Fig 5B-5D**). The CV026 strain showed significant growth inhibition at 8 μg/mL, 4 μg/mL, and 2 μg/mL for CT **(Supplementary Fig 5B**). CF also showed a similar pattern of inhibition at 4 μg/mL, 2 μg/mL, and 1 μg/mL concentrations **(Supplementary Fig 5C**). Similarly, CP inhibited growth at 64, 32, 16 and 8 μg/mL, however there was no significant effect on growth at 4 μg/mL, 2 μg/mL, 1 μg/mL, and 0.5 μg/mL concentrations **(Supplementary Fig 5D**). To quantify the anti-QS activity using pigment inhibition, we selected the 1/2^th^, 1/4^th^, 1/8^th^, and 1/16^th^ concentrations of MIC values of CP, CT, and CF. Exogenous AHL was supplied in liquid media. After overnight incubation, CV026 showed purple pigment production and microtiter wells appeared purple. Quantification of the pigment was performed by measuring the OD at 567 nm. The intensity of each well was compared with the control. CP at 32.0 μg/mL and 16.0 μg/mL significantly (p<0.05) inhibited pigment production (**Fig 1D** and **1G**), while CF inhibited pigment production even at 2.0 μg/mL (**Fig 1E** and **1H**). CT showed significant pigment inhibition at 4.0 μg/mL (**Fig 1F** and **1I**). The plate reader-based quantitative analysis confirmed that cephalosporins inhibited QS pathways in *C. violaceum* at sub– MICs of CP, CF, and CT.

### Molecular docking of cephalosporins to the CviR binding pocket

To investigate the mechanism of QS inhibition, we performed molecular docking of the CviR receptor with CP, CF, and CT using Autodock Vina (**Fig 1J-1M**). To find interacting amino acids in the binding pockets of the QS receptor CviR with the ligand, we selected amino acids within 5Å distance of each ligand. The natural ligand of the CviR receptor, C10-HSL, showed the highest predicted binding affinity with −6.9 kcal mol^-1^. The stabilization of C10-HSL in the binding pocket of the receptor is based on hydrogen bond interactions of the carbonyl group with Trp84, the amide carbonyl group with both Tyr80 and Ser155, and the secondary amine with Asp97 **(Fig 1J)**. The last carbon on the alkane tail of C10-HSL formed an alkyl interaction with Met72 and a π-alkyl interaction with Tyr88. Of the inhibitors, CT displayed the most favorable interactions with the CviR receptor, with a predicted affinity of −6.5 kcal mol^-1^. CT formed four hydrogen bonds; one between Met72 and the carbonyl on the beta-lactam ring, one between Ile69 and the carboxylic acid, and two more hydrogen bonds with Tyr88 and Asn92 from the amine connected to the aromatic ring **(Fig 1M)**. A π-sulfur interaction was formed between Met72 and the aromatic ring, an alkyl interaction between Ala94 and the non-aromatic ring thioether, as well as a π-alkyl interaction with the aromatic ring Ala94. In addition to this, an unfavorable interaction was formed between Ala94 and the proton of the amine connected to the aromatic ring, which may be the reason for the slightly lower predicted energy than the natural ligand.

CP showed the second highest predicted binding affinity of −6.3 kcal mol^-1^ with the CviR receptor. CP formed π-alkyl interactions with Met72 and Ala94 and an amide-π stacking interaction with Gln70 **(Fig 1K)**. Two hydrogen bonds were formed between Ile69 and the proton of the primary amine connected to the aromatic ring, and between Met72 and the nitrogen in the methoxy-amino group. Alkyl interactions were formed with Met72 and Val75 residues. In addition, an unfavorable donor-donor interaction was detected between the proton of the carboxylic acid and Ser89.

Among the three inhibitors, CF showed the lowest affinity for the CviR receptor (−6.1 kcal mol^-1^). CF formed a hydrogen bond with Ser89 from the primary amine connected to the aromatic ring, and another with Met72 from a carboxylic acid group. Near the carboxylic acid group, four alkyl interactions were formed with the neighboring carbon atoms **(Fig 1L)**.

### Effect of cephalosporins on the growth profile of *P. aeruginosa* PAO1

To estimate anti-virulence and antibiofilm activities of the selected cephalosporin antibiotics, we studied the effect of sub-MICs of CP, CF, and CT on the growth of *P. aeruginosa* PAO1. Most of the cephalosporin antibiotics showed potent antimicrobial activity against PAO1 except oxacillin (no antimicrobial activity at 265 μg/mL). MIC values for imipenem, meropenem, doripenem, ertapenem, CT, CF and CP were found to be 256 μg/mL, 2 μg/mL, 32 μg/mL, 1 μg/mL, 32 μg/mL, 4 μg/mL, and 2 μg/mL (**Supplementary Fig 6**). Interestingly, the OD600 values at sub-MIC of each antibiotic were found to be significantly lower (p<0.001) than the control group (PAO1) (**Supplementary Fig 7A-7H**). This indicated that growth was influenced by antibiotics. Surprisingly the growth was also significantly affected in oxacillin antibiotics that showed no anti-microbial activity against PAO1 (512-0.5 μg/l) (**Supplementary Fig 6E**). These results were encouraging and indicated that antibiotics were targeting alternate molecular pathways influencing the growth of *P. aeruginosa*. Further, we measured growth curves of PAO1 in the presence and the absence of sub-MIC (MIC, MIC/2, and MIC/4) of each antibiotic (**Fig 2A-2C**). We also compared the growth of PAO1 with QS mutant strains as shown in **Fig 2D**.

**Figure-2:**
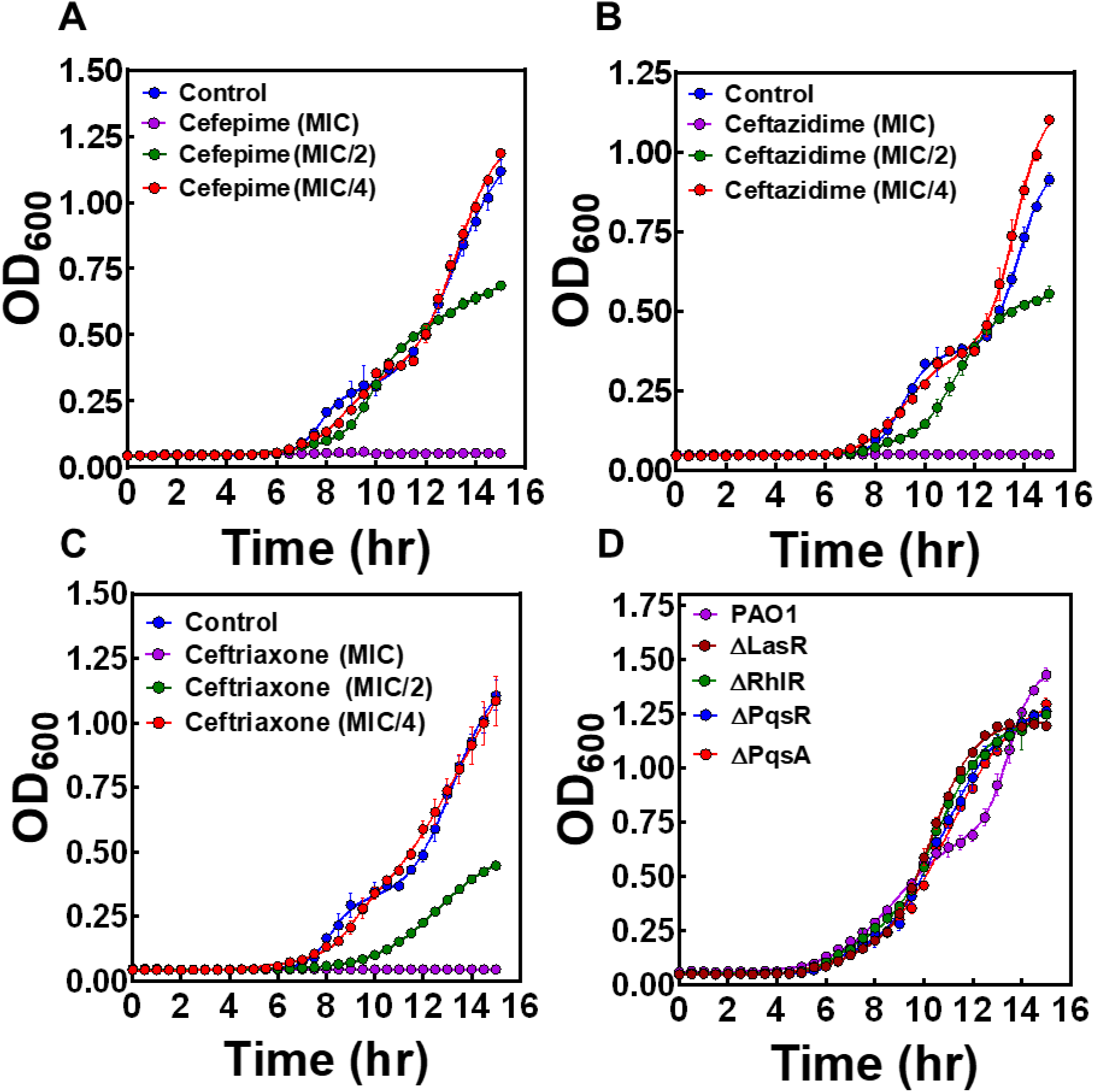
Effects of cephalosporins on the biphasic growth of *P. aeruginosa* PAO1. Growth curves of PAO1 at different concentrations (MIC, MIC/2, and MIC/4) of CP (A), CF (B), and CT (C) respectively. Growth curves of *P. aeruginosa* PAO1 and its isogenic QS-mutant strains (ΔLasR, ΔRhlR, ΔPqsA, and ΔPqsR) (D).

### Effect of cephalosporins on pyocyanin production

In the present study, the antibiotic treatment showed significant inhibition of pyocyanin production at a sub-MIC concentration as compared to control (**Fig 3A** and **3B**). PAO1 showed high pyocyanin pigment production OD690 nm (1.67 + 0.19) and with CT (1.02 + 0.18), CF (0.77 + 0.27), CP (0.48 + 0.17), pyocyanin production was significantly reduced (**Fig 3A** and **3B**). These results collectively suggested that CP, CF, and CT interfered with QS pathways leading to the inhibition of pyocyanin production. As shown in **Fig 3**, there is a clear reduction of pyocyanin production by CP, CF, and CT.

**Figure-3:**
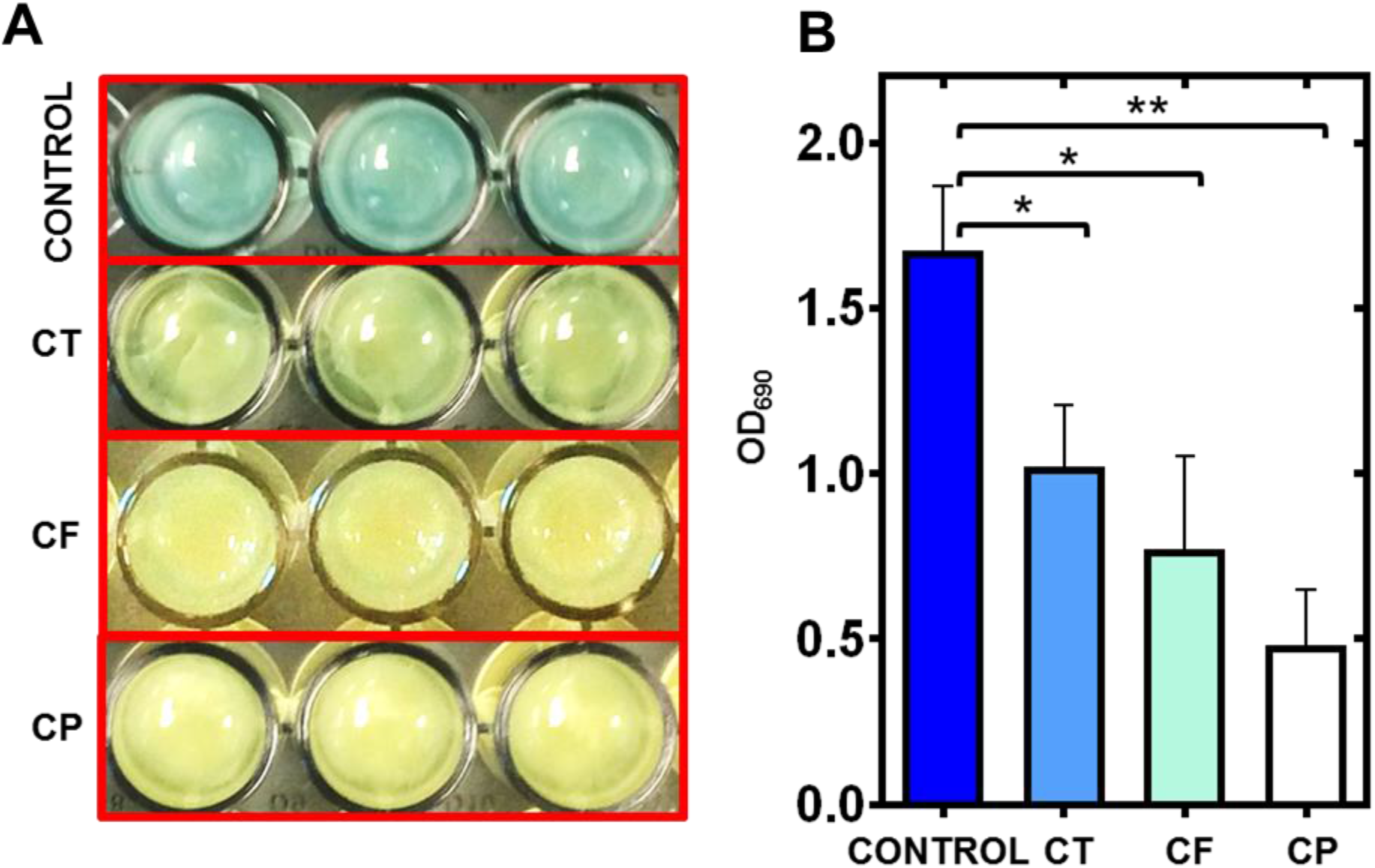
Inhibition of pyocyanin production of *P. aeruginosa* PAO1 by cephalosporins. Image of microtiter plate wells (A) and graphical representation of OD690nm (B) showing the inhibition of blue color (pyocyanin production) by *P. aeruginosa* PAO1 in presence of CT (CT), CF (CF) and CP (CP) (* p ≤ 0.05 and ** p ≤ 0.01).

### Effect of cephalosporins on motility phenotypes of *P. aeruginosa* PAO1

We have tested the motility phenotypes in the presence of sub-MIC of selected antibiotics. Swimming motility was analyzed for control and QS mutant strains. We found that the QS mutant strains showed no significant difference in swimming motility, which suggests that the flagella-dependent swimming motility is independent of QS pathways (**Fig 4A_i_** and **Fig 5A**). Interestingly all three antibiotics CP, CF, and CT showed significant swimming motility inhibition (p<0.001) of PAO1 as well as the isogenic QS mutant strains (**Fig 5A**). Diameters of the zones are mentioned as table insert in **Fig 5A**. Further, the effect of antibiotics on swarming motility was also analyzed and it was found that the ΔLasR, ΔRhlR mutant strains showed significant difference (p<0.05) as compared to the control indicating that the motility is dependent of LasR and RhlR QS pathways (**Fig 4B_i_** and **Fig 5B**). PqsR and PqsA mutant strains showed no significant difference in the swarming motility zone as compared to the control PAO1. Interestingly all three antibiotics CP, CF, and CT showed significant swarming motility inhibition (p<0.001) for all strains (**Fig 4B_i_** and **Fig 5B**). Effect on twitching motility was also analyzed on PAO1 and its isogenic mutant strains, and it was found that the ΔLasR, ΔRhlR mutant strains showed significant difference (p<0.05) in the twitching motility (**Fig 4C_i_** and **Fig 5C**) indicating that the twitching motility is dependent on las and rhl QS pathways. Interestingly only two antibiotics CP and CT showed significant twitching motility inhibition (p<0.001) but surprisingly CF showed no motility inhibition (**Fig 4C_i_** and **Fig 5C**). Our results clearly showed that CP and CF have a strong inhibitory effect on twitching motility indicating their potential effect on the establishment of infection and biofilm formation by *P. aerugionsa*. Motility inhibition activity of cephalosporins has encouraged us to evaluate the antibiofilm activity of cephalosporins against PAO1.

**Figure-4:**
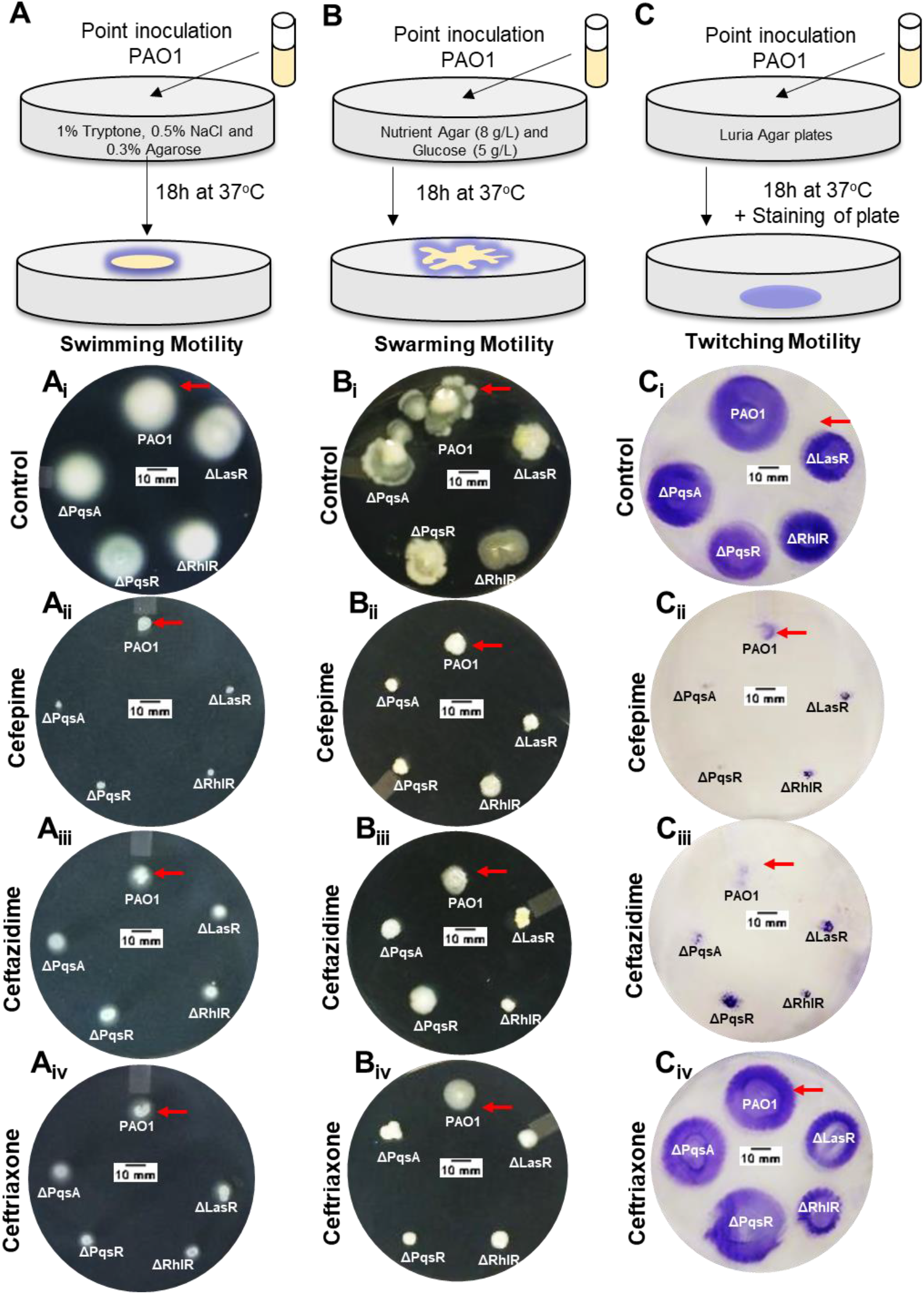
Effect of cephalosporins on motility phenotypes of *P. aeruginosa* PAO1 and its isogenic QS mutant strains. Image showing the comparative zone of swimming (A, A_i_), swarming (B, B_i_), and twitching motility (C, C_i_) of PAO1 and its isogenic QS mutant strains (ΔLasR, ΔRhlR, ΔPqsA, and ΔPqsR). Image of the media plate showing the effect of CP (A_ii_, B_ii_, and C_ii_), CF (A_iii_, B_iii_, and C_iii_), and CT (A_iv_, B_iv_, and C_iv_), on swimming (A), swarming (B) and twitching motility (C) zones of *P. aeruginosa* PAO1 (Scale bar=10 mm).

**Figure-5:**
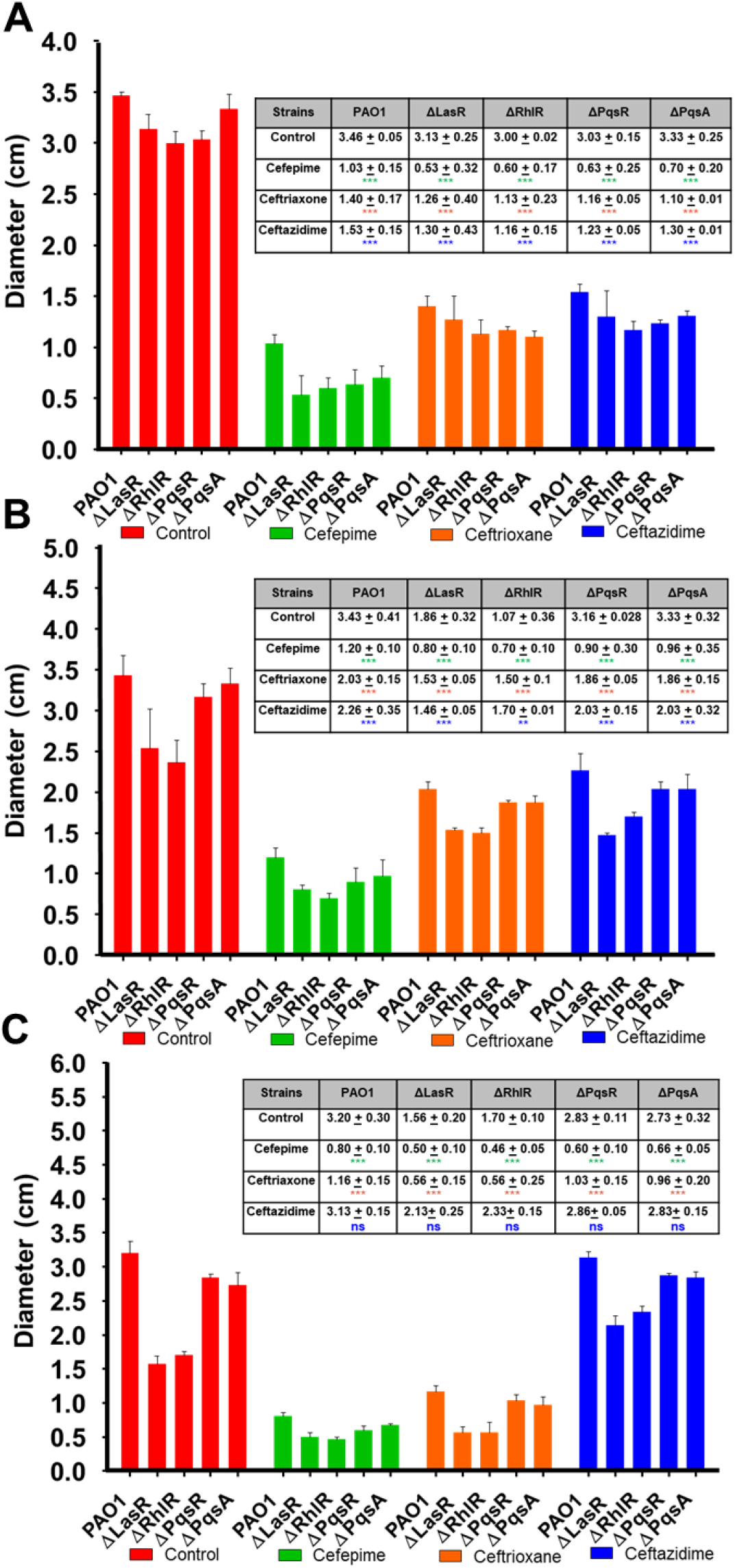
Quantification of motility inhibition (diameter of motility zones) of *P. aeruginosa* by cephalosporins: Comparative analysis of the diameter of the motility zone (mm) for PAO1, ΔLasR, ΔRhlR, ΔPqsR and ΔPqsA in presence of CP (A), CF (B) and CT (C). Inset table showing the diameter of zone (mm) of swimming, swarming, and twitching motility phenotypes for PAO1 and its isogenic QS mutant strains. Not significant (ns): p > 0.05, * p ≤ 0.05, ** p ≤ 0.01, *** p ≤ 0.001, **** p ≤ 0.0001.

### Anti-biofilm activity of cephalosporins against *P. aeruginosa* PAO1

To test whether QS inhibition by cephalosporins (sub-MICs that do not influence growth profile) also affects biofilm formation by *P. aeruginosa*, we used Environmental Scanning Electron Microscopic (ESEM) to image biofilms produced by PAO1 (**Fig 6**). We have tested the effect of antibiotic supplementation from day 1 to day 7. SEM imaging and live cell counts were performed on every alternate day (days 1, 3, 5, and 7) on the urinary catheter surface. The ESEM technique allowed us to visualize the biofilm under their natural state without any physical or chemical treatment (no dehydration and staining). On day 1, PAO1 control biofilms showed the presence of a uniform monolayer of cells attached to the urinary catheter surface. Cells were uniformly distributed within the extracellular polysaccharide matrix (**Fig 6A_i_**). The live cell count of the day 1 biofilm was high (6.39±1.89 log cfu) (**Fig 6A**) indicating the PAO1 cells successfully attached to the catheter surface at high cell numbers. Interestingly, even in antibiotic-treated groups, there was a small number of cells attached to the catheter surface. CP showed low cell counts with significantly less polysaccharide matrix distributed throughout the catheter surface indicating the inability of *P. aeruginosa* cells to attach to the catheter surface (**Fig 6A_ii_**). CP treatment significantly reduced (p<0.001) the log cfu count as compared to the control (3.83±0.32 log cfu). Interestingly, CF and CT both showed a significant amount of exopolysaccharide distributed on the catheter surface but minimum cell attachment (**Fig 6A_iii_** and **A_iv_**). Both antibiotics showed reduced live cell counts of 4.70±0.26 and 5.50±0.43 log cfu (**Fig 6A**). This indicated that the cells were defective in the production of exopolysaccharide and were unable to attach to the catheter surface. Three-day-old-biofilms of PAO1 of the control group showed the formation of a thick layer of bacterial cells embedded in an exopolysaccharide matrix (**Fig 6B_i_**). Biofilms appeared to be complex in structure with the presence of long polymer threads, mucilaginous matrix, and high cellular growth. Live cell counts were higher on day 3 in control groups (8.28±1.13 log cfu) (**Fig 6B**). The CP-treated catheter surface showed low cell counts attached to the urinary catheter surface at day 3 indicating the potent anti-adherence ability of CP (**Fig 6B_ii_**). The bacterial live cell count on day 3 CP treated urinary catheters was lower (5.01 + 0.56 log cfu) as compared to the control live cell count (**Fig 6B**). CF and CT antibiotics showed thin biofilm formation on the catheter surface (**Fig 6B_iii_** and **B_iv_**). The bacterial cells appeared to be longer in size as compared to the control. These results indicate that the effect of antibiotic treatment altered the cell physiology and growth cycle of *P. aeruginosa*. Log cfu count of the treated groups were significantly reduced (p<0.05) as compared to the PAO1 control (CT 6.0±0.56 log cfu and CF 6.6±0.73 log cfu) (**Fig 6B**). Five-day-old-biofilms of the PAO1 control group showed thick biofilm and completely covered with a thick exopolysaccharide layer (**Fig 6C_i_**). On day 5, the biofilms started to construct mushroom-shaped structures. At this stage, biofilms might be completely resistant to antimicrobial agents due to the protection provided by the exopolysaccharide layer. These clusters of live bacterial cells help in the formation of the three-dimensional architecture of bacterial biofilm. As expected, the biofilm showed a high live cell count in five-day-old biofilms (10.05±0.86 log cfu) (**Fig 6C**). In CP treated groups, the biofilm appeared to be defective (**Fig 6C_ii_**) and the log cfu count of the live cell was significantly lower indicating the potent antibiofilm activity of CP (**Fig 6C**). In CT and CF, treated biofilms were condensed and covered with thick polysaccharide matrix on the catheter surface (**Fig 6C_iii_** and **C_iv_**). In CF treated biofilms, bacterial cells appeared long chains embedded in the polysaccharide matrix. However, there was no significant difference in the log cell count of live cells in CF treated biofilms as compared to the control group. CT treated biofilms showed a significant difference in the log cfu count of bacterial cells (7.01±0.19 log cfu) (**Fig 6C**). Moreover, on day-7 control biofilms appeared highly complex three dimensional with numerous mushroom-shaped secondary structure on the biofilm surface (**Fig 6D_i_**). In the untreated control group, biofilms showed a larger assembly of multi-mushroom shaped structures, and the log cfu count of live cells in 7-day-old-biofilm was found to be very high (12.60±0.75 log cfu) (**Fig 1D**). The seven-day-old CP treated biofilms were still defective in biofilm architecture. The mushroom-shaped structures were absent and individual cells can be visualized on the catheter surface. In treated biofilms, bacterial cells appeared as long thread shaped and produced significantly less polysaccharide with low log cfu count of 7.81±0.65 (**Fig 6D**). CT and CF biofilms were covered with mushroom-shaped structures and few individual cells were visible indicating the thick threedimensional biofilm formation. Live cell count of the catheter surface of these antibiotic-treated groups was found to be non-significant as compare to control indicating the cells have counteracted the effect of these antibiotics using complex exopolysaccharide shield (**Fig 6D**).

**Figure-6:**
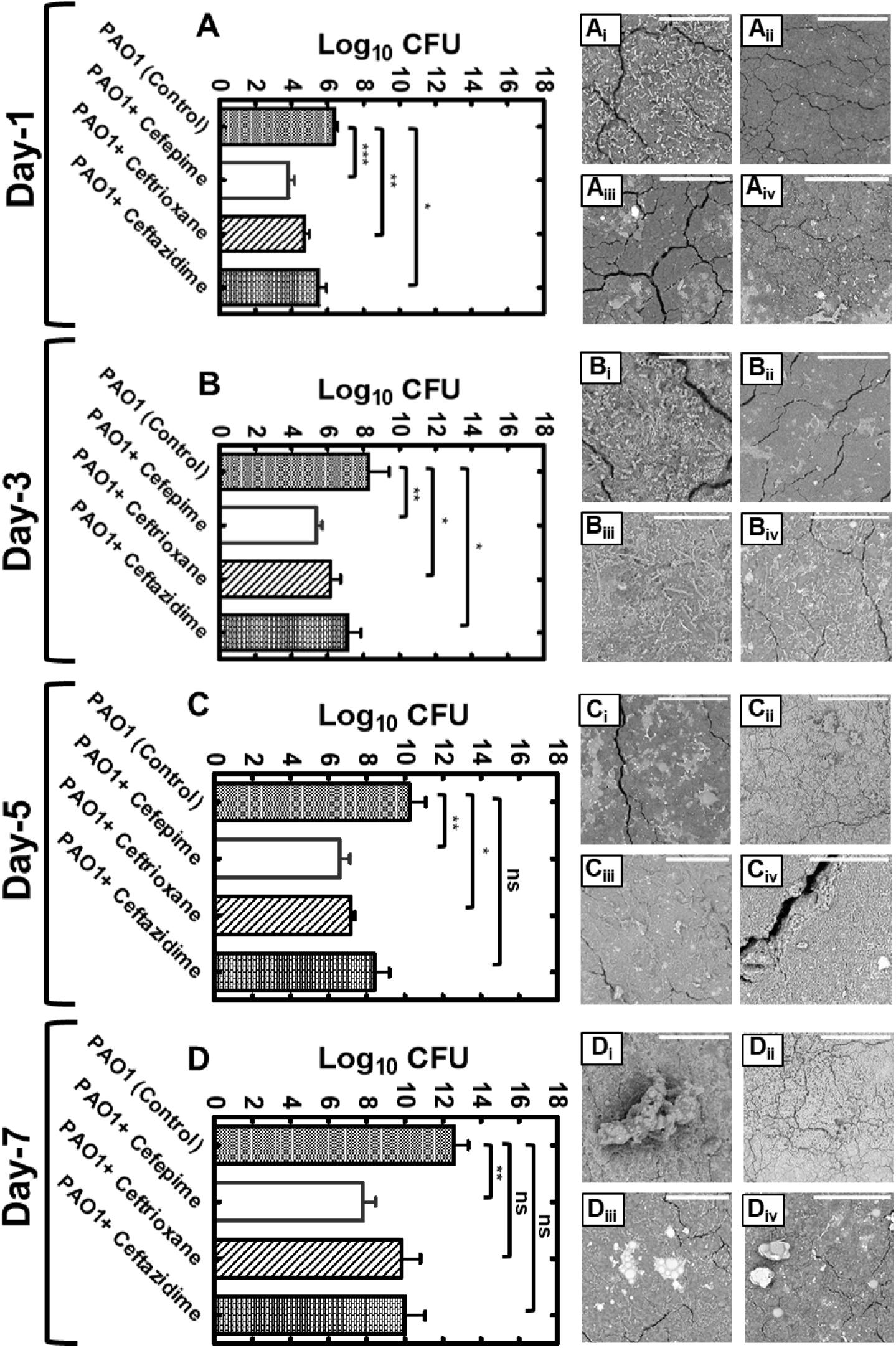
Effects of QS inhibition by cephalosporins on biofilm formation by PAO1. Graphical representation of live bacterial cell quantification of *P. aeruginosa* PAO1 on urinary catheter surface under sub-MIC concentrations of CP, CT and CF for day-1 (A), day-3 (B), day-5 (C) and day-7 (D) (ns p > 0.05, * p ≤ 0.05, ** p ≤ 0.01, *** p ≤ 0.001). Scanning Electron Microscope (SEM) images of biofilms formed by *P. aeruginosa* PAO1 on the urinary catheter in presence of CP, CF, CT on day-1 (A_ii_, B_ii_, C_ii_, and D_ii_), day-2 (A_ii_, B_ii_, C_ii_, and D_ii_), day-3 (A_iii_, B_iii_, C_iii_, and D_iii_) and day-4 (A_iv_, B_iv_, C_iv_, and D_iv_) (Scale bars represent 50 μm).

### Synergistic effect of cefepime and aminoglycosides against *P. aeruginosa* PAO1

After determining the anti-QS and anti-virulence effect of cephalosporin antibiotics against PAO1, we were interested in testing the synergistic activity of CP with aminoglycosides against *P. aeruginosa* PAO1. First, the antimicrobial efficacy of aminoglycosides was tested against *P. aeruginosa* PAO1. Kanamycin sulfate showed poor antibacterial activity against *P. aeruginosa* PAO1 (MIC: 64 μg/mL), however, in presence of CP, the MIC of kanamycin has significantly decreased to 16 μg/mL (**Fig 7A**). Streptomycin sulfate showed potent efficacy against PAO1 and the MIC was found to be 8 μg/mL (**Fig 7B**). In synergy with CP, the MIC of streptomycin was significantly reduced to 2 μg/mL. Similarly, the MIC of neomycin was found to be 16 μg/mL and in combination with CP, the MIC was decreased to 8 μg/mL (**Fig 7C**). In addition, gentamicin and tobramycin were found to be highly effective antibiotics against PAO1 with the MIC values of 2 μg/mL (**Fig 7D** and **7E**). However, CP also showed a synergistic effect with these antibiotics and the MIC values were decreased to 1 μg/mL. In literature, the anti-QS compound has shown to potential the anti-microbial activities of various antibiotics. The anti-QS compounds target various pathways that silence the expression of virulence factors. Major virulence factors including lipopolysaccharide, alginate, and extracellular matrix proteins help in the development of antimicrobial resistance and blocking QS pathways may lead to increased susceptibility of antimicrobial drugs due to suppressed virulence state of the pathogen. Our results showed that the CP at a sub-inhibitory concentration in combination with major aminoglycosides led to the increased antimicrobial potency of aminoglycosides due to its potent anti-virulence activity. This combination needs to be tested in animal models to validate its effectiveness against *P. aeruginosa* and could be established as a clinically useful combinational therapy to treat biofilm-associated *P. aeruginosa* infections.

**Figure-7:**
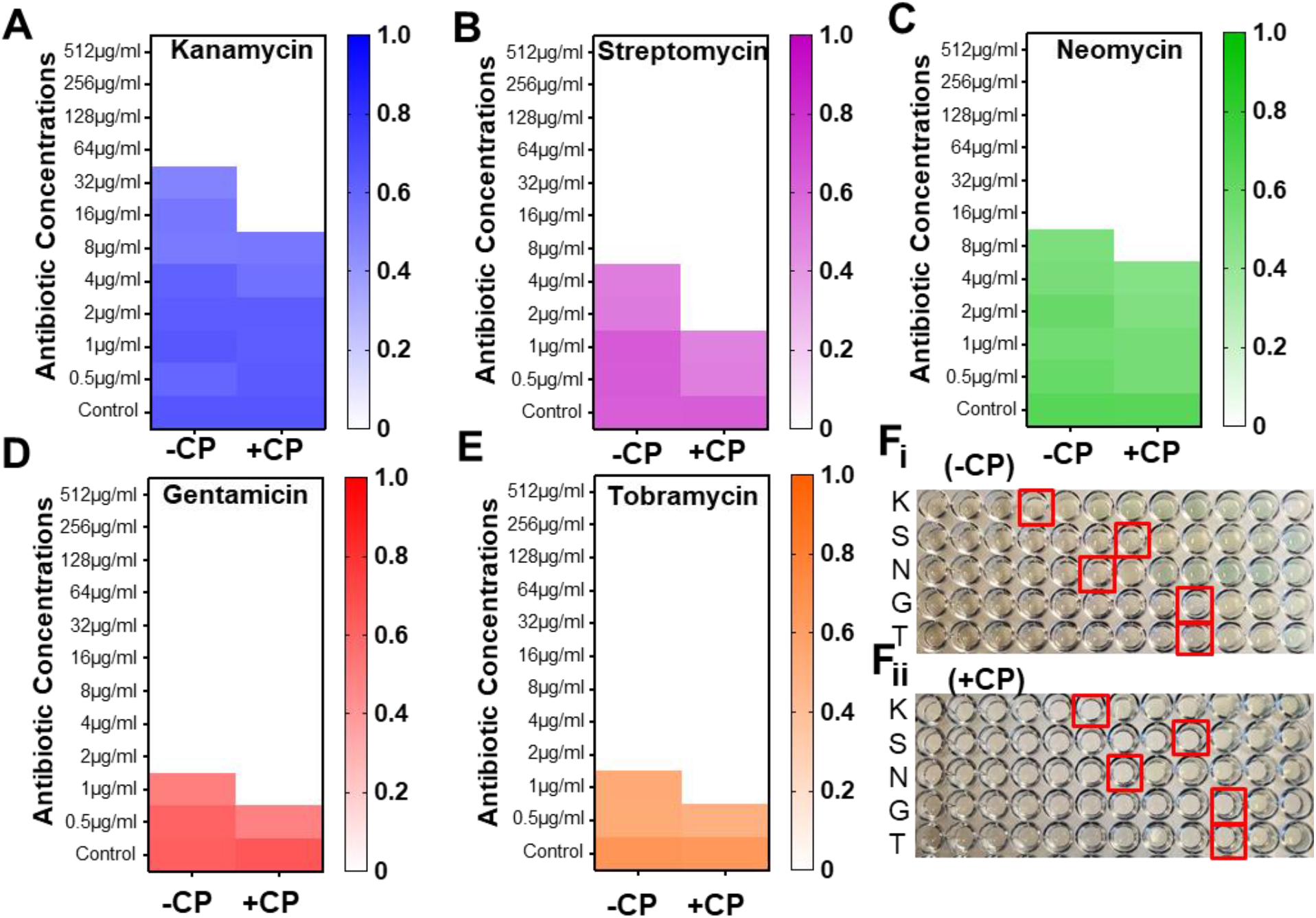
Synergistic antimicrobial effects of CP and aminoglycosides against *P. aeruginosa* PAO1. Heat map of OD600nm of *P. aeruginosa* PAO1 with kanamycin (A), streptomycin (B), Neomycin (C), Gentamicin (D), and Tobramycin (E) in presence and absence of CP. Image of microtiter plate showing the MIC of aminoglycosides against *P. aeruginosa* PAO1 in the 96-microtiter plate with and without CP (F_i_, F_ii_). (Red rectangular boxes in the image represents the MIC of aminoglycosides against PAO1-wells with no visible bacterial growth); K-kanamycin, S-streptomycin, N-Neomycin, G-Gentamicin, and T-Tobramycin.

### Molecular docking of cephalosporins with *P. aeruginosa* quorum-sensing receptors

To compare cephalosporin binding to the LasR interactions with the natural ligand, we first docked 3-oxo-C12HSL to its binding pocket in the LasR receptor. The predicted binding affinity was −8.0 kcal mol^-1^. 3-oxo-C12HSL formed three hydrogen bonds, one with Ser129 and the amide carbonyl, and two more with Asp73, Thr75, and the proton of the secondary amine (**Fig 8A**). A hydrophobic interaction connected Tyr56 and the ring carbon neighboring the secondary amine, supported by two alkyl interactions between the carbon at the end of the hydrophobic tail with Leu40 and Ala50. Next, the three selected antibiotics were docked to the same ligand-binding pocket of LasR. CP formed two hydrogen bonds, one between the carboxyl group and Asp65 and another between Ala50 and the proton of the secondary amine connected to the beta-lactam ring (**Fig 8B**). A π-sulfur interaction was formed between Phe167 and the sulfur atom of the aromatic ring, an electrostatic interaction was formed between Asp65 and the charged nitrogen atom, a carbon-hydrogen bond was formed between Asn49 and the beta-lactam carbonyl, and an alkyl interaction was formed between Ile52 and the thioether group. CP showed a comparative docking score of - 5.6 kcal mol^-1^. CF formed three hydrogen bonds; one between Gly54 and a carboxyl group, and two between the primary amine with Tyr56 and Asn55. Lys16 formed a carbon-hydrogen bond with the carboxyl group and an alkyl interaction with a nearby methyl group (**Fig 8C**). A pi-alkyl interaction was formed between Ala58 and the aromatic ring, and an unfavorable electrostatic interaction was formed between Glu62 and the carboxyl nearest to the beta-lactam ring. CF showed a comparatively low docking score of −5.9 kcal mol^-1^. CT formed two hydrogen bonds; one between Asn49 and the primary amine and another between Lys16 and the non-aromatic ring thioether (**Fig 8D**). A pi-sulfur interaction was formed between Phe167 and the aromatic ring thioether. Carbon hydrogen bonds were formed between Ala58 and the methyl group connected to the triazine ring, and between Asn49 and the beta-lactam carbonyl. Three alkyl interactions were formed; one from Ile52 to the non-aromatic thioether ring, two from the triazine ring to Ala58 and Arg61, respectively. CT showed a comparatively higher docking score of −6.6 kcal mol^-1^.

**Figure-8:**
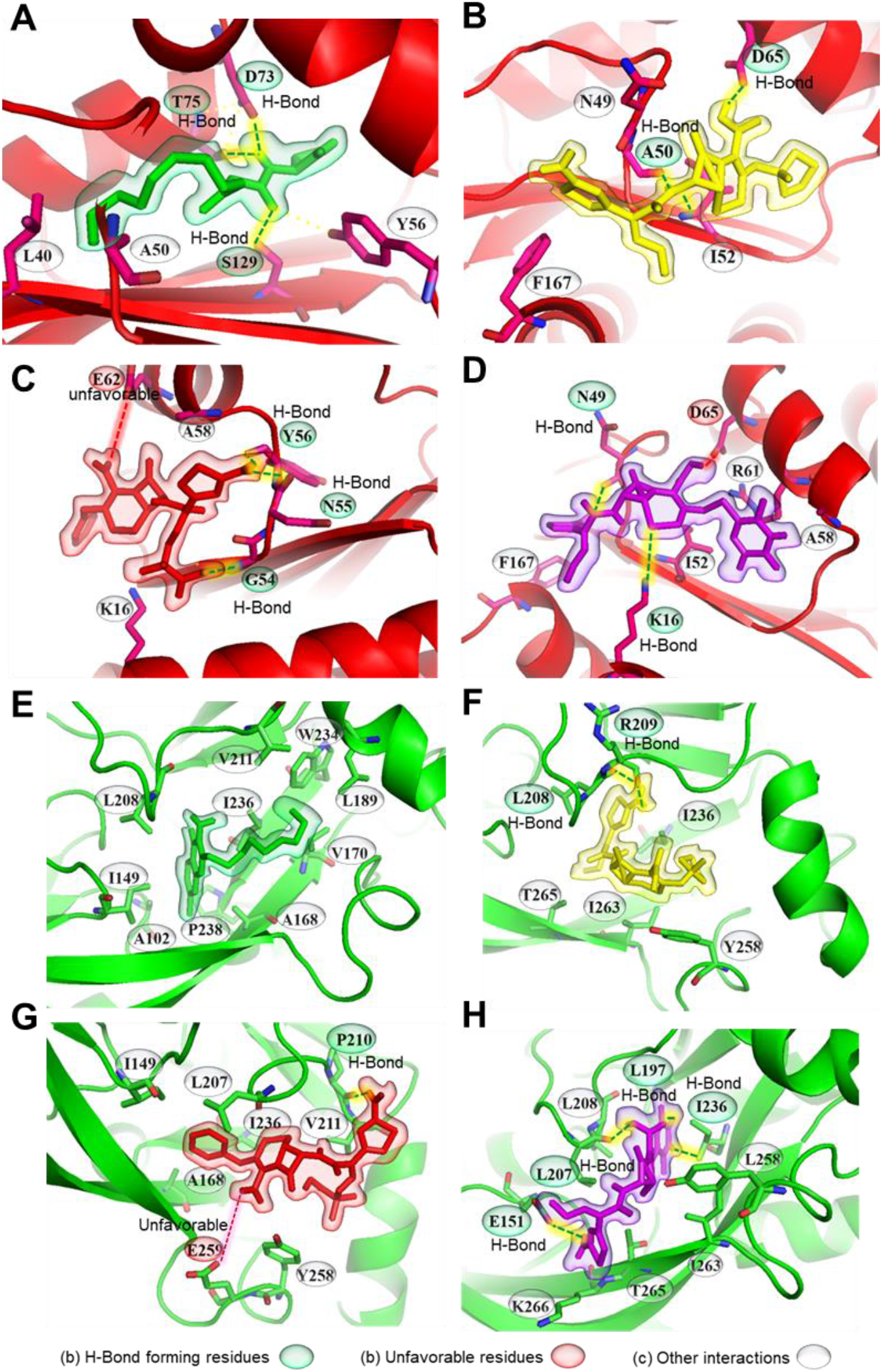
Molecular interactions of natural ligand and cephalosporin antibiotics with *P. aeruginosa* QS receptors (LasR and PqsR). Representation of ligand-receptor interactions of 3oxo-C12-HSL (A), CP (B), CF (C), CT (D) with LasR QS receptor. Representation of ligand-receptor interactions of Pseudomonas quinolone signaling (PQS) molecule (E), CP (F), CF (G), CT (H) with PqsR QS receptor. LasR and PqsR receptors are represented as red and green. Natural ligands of receptors, CP, CF, and CT are presented as green, yellow, red, and purple.

Molecular docking was also performed independently with the PqsR receptor. Interestingly, the natural ligand formed no hydrogen bonds but formed ten alkyl interactions between its two rings with Leu208, Ala168, Ile149, Pro238, and Ala102, as well as five alkyl interactions between the last carbon in the hydrophobic tail and Leu189, Val170, Trp234, Val211, Ile236 of PqsR receptor (**Fig 8E**). The natural ligand showed a docking score of −7.4 kcal mol^-1^. CP formed two hydrogen bonds from its primary amine to Arg209 and Leu208. A carbon-hydrogen bond was formed between the methoxyl group and Thr265 (**Fig 8F**). Two π-σ interactions were formed from the aromatic ring to Leu208 and Ile236. An alkyl interaction and a π-alkyl interaction were formed from the thioether to Ile263 and Tyr258, respectively. The docking score of CP was found to be - 6.7 kcal mol^-1^. Similarly, CF formed a hydrogen bond between its primary amine and Pro210 as well as a carbon-hydrogen bond from Leu207 to the six-membered ring (**Fig 8G**). An unfavorable interaction was also formed between Glu259 and the carboxyl nearest the beta-lactam ring. CF showed the docking score of −6.2 kcal mol^-1^. CT formed four hydrogen bonds; one between its primary amine and Glu151, one between Leu207 and the carboxyl group, one between Ile236 and the proton on the triazine ring, and lastly one between Leu197 and a carboxyl on the triazine ring (**Fig 8H**). The docking score of CT was −6.7 kcal mol^-1^. These conformations would obstruct the entry into the binding pocket, and therefore interrupt ligand-receptor interactions. Thus, the molecular docking analysis results indicated that the cephalosporin antibiotics suppress QS signaling of *P. aeruginosa* by competitive inhibition of LasR.

### Protective effect of cephalosporins in the *C. elegans* infection model

We have used *C. elegans* as an animal model to investigate the anti-virulence effect of CP, CF, and CT against *P. aeruginosa* PAO1. Negative control (untreated) *C. elegans* were found to be healthy and motile. Advantage of its transparent nature, internal organs can be visualized directly and in healthy worms, the intestine, uterus, proximal and distal gonads (Fig 9A_i_ and A_ii_) were found to be intact and without inflammation [47]. The dead worm has blunt-body curvature with reduced body movement; these were distinct phenotypic characteristics of the inflammatory or diseased state of the *C. elegans*. PAO1 infected *C. elegans* showed potent inflammatory damage of intestinal tissue and tissue necrosis after 72 hr leading to 100% mortality in infected worms [47]. Percentage survival of *C. elegans* at 12, 24 and 28 h was 88.0 + 3.60%, 65.0 + 4.30% and 36.6 + 8.3 0% and at 72 h, it has achieved 0% survival. All worms were found dead after 72 h (Fig 9F). Interestingly, the sub-inhibitory concentration of antibiotic treatment showed reduced virulence in *C. elegans*. CP treatment showed a significant increase in the percentage survival of *C. elegans*. At 12, 24, 48 and 72h the % survival was 95.6 + 2.51%, 92.33 + 2.50%, 83.33 + 4.16%, and 65.33 + 5.03% (Fig 9F). Similarly, CF treated supernatant also showed a significant increase in worm survival. At 12, 24, 48 and 72 h the % survival was found to be 93.6 + 1.52%, 85.33 + 5.03%, 77.0 + 5.56% and 58.0 + 4.30% (Fig 9F). CT showed similar anti-virulence effect and treatment groups showed reduced mortality. At 12, 24, 48 and 72 h, the percentage survival was found to be 92.0 + 2.60%, 84.33 + 5.56%, and 48.66 + 8.08% (Fig 9F). All three antibiotics showed a significant reduction in the *C. elegans* mortality indicating the potent antivirulence effect of a sub-inhibitory concentration of cephalosporin antibiotics. *C. elegans* was found to be healthy and inflammatory damage of intestinal tissue was significantly reduced indicated the strong antivirulence effect of cephalosporins. Subsequently, to prove antivirulence activity is related to QS inhibition of *P. aeruginosa*, we tested this hypothesis by evaluating the effect of QS mutant strains of *P. aeruginosa* on *C. elegans* survival. Results showed that isogenic QS mutant strains were defective in virulence against *C. elegans*. Percentage survival of ΔLasR mutant supernatant at 12, 24, 48 and 72 h was 92.66 + 1.52%, 82.0 + 5.29%, 59.0 + 9.53% and 31.66 + 11.23% (Fig 9G). Similarly, ΔRhlR mutant strains showed 89.0 + 3.60%, 77.6 + 10.20%, 55.0 + 10.0% and 27.33 + 7.63% (Fig 9G). Similarly, ΔPqsR and ΔPqsA mutant strains were also found to have 86.33 + 5.13%, 71.33 + 6.11%, 47.66 + 11.23%, 26.66 + 10.40% and 84.33 + 4.04%, 75.66 + 7.76%, 57.0 + 12.53%, 36.66 + 5.77% survival rates of *C. elegans* (Fig 9G). We have also used an uninfected group with *E.coli* OP50 to check the *C. elegans* mortality and no mortality was observed until 72 hr. The result of our experiment showed that the sub-MIC concentration of antibiotics suppressed the virulence factors of *P. aeruginosa* and led to an increase in the survival of *C. elegans*. Finally, we have compared the percentage mortality of all the groups with the control PAO1 group and all the groups have shown a significant reduction in the percentage mortality at 72 hr (Fig 9H). Additionally, mortality rates were higher in the QS mutant strains as compared to the antibiotic-treated groups (Fig 9G). This indicates that the single QS mutation does not completely makes the bacteria avirulent, therefore, the therapies targeting multiple QS systems will be useful as antivirulence drugs. This also indicates that cephalosporins might have additional alternative antivirulence mechanism apart from anti-QS activity that may target virulence of *P. aeruginosa*. In the future, it might be interesting to investigate the antivirulence studies at the molecular level and validate the CP, CF, and CT as potential antivirulence therapies to treat *P. aeruginosa* infections.

**Figure-9:**
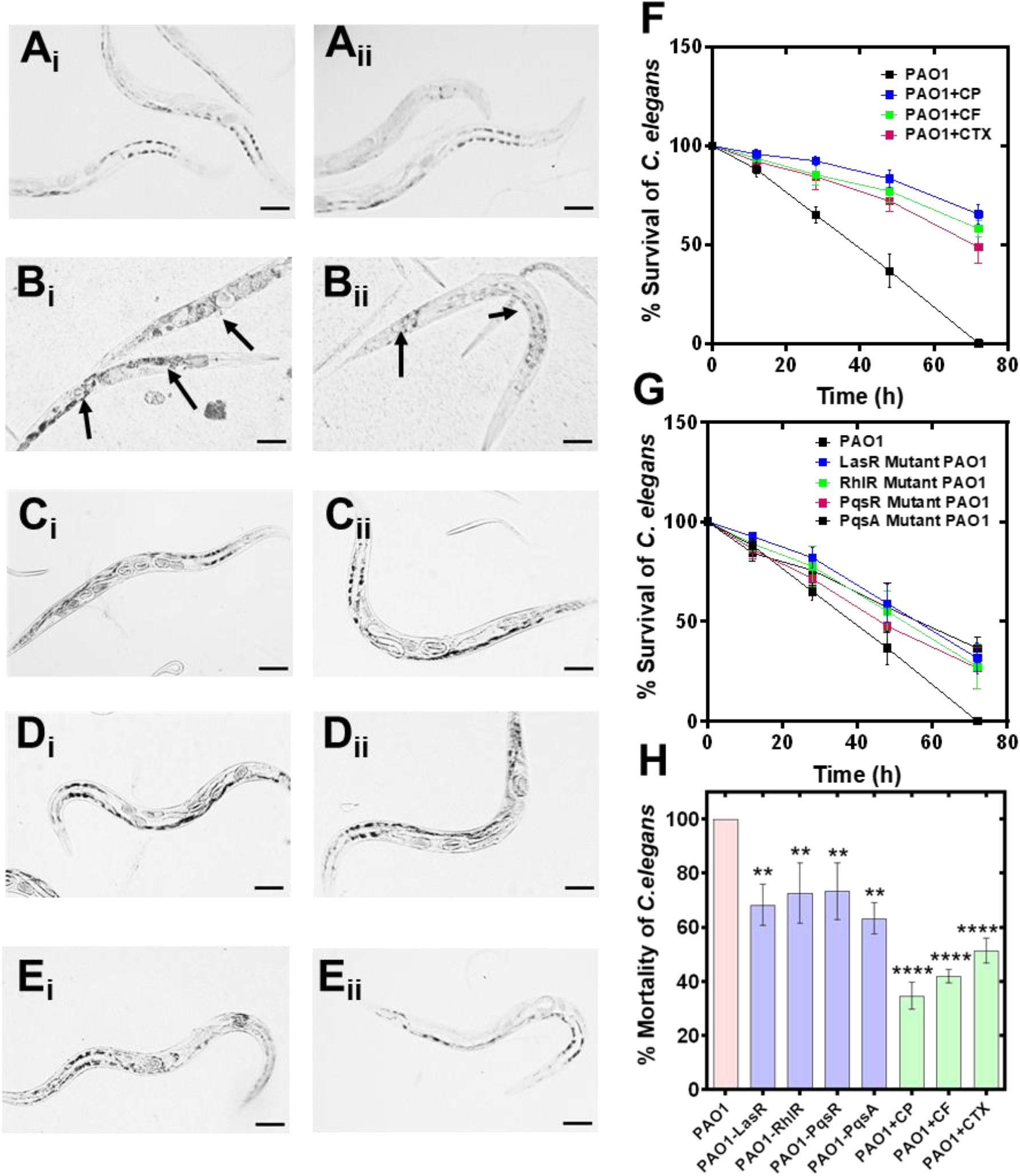
Antivirulence effect of cephalosporins against *P. aeruginosa* PAO1 induced inflammatory damage in *C. elegans*. Light microscopic images of *C. elegans* showing the protective effect of cephalosporin in *P. aeruginosa* mediated inflammatory; control healthy *C. elegans* (A_i_, A_ii_); *C. elegans* exposed to PAO1 (B_i_, B_ii_); *C. elegans* exposed to PAO1 in presence of CP (C_i_, C_ii_); CF (D_i_, D_ii_), and CT (E_i_, E_ii_). Percentage survival of PAO1 infected *C. elegans* in the presence of CP, CF, and CT (F). Percentage survival of *C. elegans* by the QS mutant strains (ΔLasR, ΔRhlR, ΔPqsR, and ΔPqsA) and control PAO1 strain (G). Comparative analysis of the % mortality of *C. elegans* after 72 hr by control PAO1 strain, isogenic QS mutant strains (ΔLasR, ΔRhlR, ΔPqsR, and ΔPqsA) and after treatment with CP, CF, and CT (H).

## DISCUSSION

### Cephalosporins showed potent antimicrobial activity and interfered with the QS pathway of *C. violaceum* at sub-inhibitory concentrations

*C. violaceum* CV026 is a Gram-negative rod-shaped (bacillus) and motile bacterium that lives in soil and water [48]. This bacterium has been named as Chromobacterium since it produces purple colored colonies on media plates [49]. The purple pigment (violacein) is secreted in response to acyl-homo-serine molecules (QS signal) in natural environmental conditions [50]. Researchers have developed its QS deficient mutant variant referred to as CV026. This strain is a transposon insertion mutant (carrying a single insertion in the AHL synthase gene) [51]. Therefore, CV026 is an AHL negative strain that does not synthesize AHL signal molecules, but this strain responds to exogenous AHL molecules (from C4 to C8 acyl side chains AHL molecules) [52]. Consequently, CV026 strain serves as a model for detecting AHL molecules and screen compounds for their anti-QS activity [53, 54]. We have utilized CV026 strain in agar well diffusion assay to screen anti-QS activity of major β-lactam antibiotics. This assay provides information on (1) anti-microbial activity and (2) anti-QS activity of tested compounds. Our results showed that all β-lactams accept oxacillin showed potent antimicrobial activity against *C. violaceum* **(Fig 1A-144 1C, Supplementary Fig 1).** Oxacillin has a narrow activity spectrum and shows potent antimicrobial activity specifically against Gram-positive bacteria including *staphylococcus aureus* [55]. Our results were in line with the previous reports suggesting that the resistance against oxacillin is common among Gram-negative bacteria [56]. The prevalence of broad-spectrum oxacillinase variants might be responsible for the oxacillin resistance among Gram-negative bacterial strains [57, 58]. β-lactam antibiotics (carbapenems and cephalosporins) target and disrupt the peptidoglycan layer of the bacterial cell wall by blocking the important transpeptidation step leading to the irreversible inhibition of penicillin-binding protein-dependent crosslinking of peptidoglycan [59]. Peptidoglycan is important for cell integrity and inhibition in the peptidoglycan biosynthesis layer induce structure instability that leads to cell lysis [60]. Structural damage of the peptidoglycan layer by β-lactams causes disintegration of the bacterial cell wall leading to bacterial cell lysis. The order of antimicrobial activity of β-lactams against C*. violaceum* was as follows (highest to lowest): MP>IP>DP>EP>CF>CT>CP>OX **(Supplementary Fig 3)**. There was a significant difference between the intensity of antimicrobial activity among carbapenems (MP, IP, DP, and EP; the average antimicrobial zone is ~5.0 mm) as compared to cephalosporins (CF, CT, and CP, average antimicrobial zone is ~2.0 mm). The variation in the antimicrobial activity of β-lactams might be due to their specific binding affinity and selectivity against peptidoglycan binding proteins and their ability to inhibit the function of PBPs. Research have shown that the binding affinity of for DP, MP, CF and CF is different for PBP1a, PBP1b, PBP2x, PBP2a, PBP2b and PBP3 [61]. The PBP binding affinity could be a potential reason for the variation in their antimicrobial activity against *C. violaceum;* however, additional parameters including bioavailability [62], lipid bilayer transport [63] and drug solubility [64] might also play a key role for their antimicrobial activity.

In a typical agar well diffusion assay, the antibiotics diffuse in media and its concentration decreases as it diffuse from wells towards the periphery. At the interface between bacterial growth and no growth, there is a zone where antibiotic is present in sub-inhibitory concentrations. The Sub-MIC is not sufficient to kill the bacteria; however, it can provide useful information on the alternative activity of antibiotics such as anti-QS activity. We analyzed our zones of sub-MIC concentrations for each antibiotic and found that the three major cephalosporins showed a large zone of pigment inhibition at the interface of growth and no growth indicating their potential anti-QS activity against *C. violaceum* CV026. The presence of the zone of pigment inhibition indicated that the cephalosporin antibiotics were interfering with the activation of the QS pathway in the CV026 strain **(Fig 1A-C).** The size of zones for the antibiotics was as follows (large to small) CP>CF>CT>EP>DP>MP>IP>OX. There was clear evidence that the carbapenem (MP, IP, DP, and EP) showed significantly low anti-QS activity as compare to cephalosporins (CP, CF, and CT). The anti-QS activity trend was completely reversed to the anti-microbial activity trend. We hypothesized the main two reasons to explain this trend (1) the small zone of anti-QS activity for carbapenem might be because of their potent anti-microbial activity as compare to the cephalosporins; (2) the potent anti-QS activity of cephalosporin as compared to the carbapenems. To quantify the anti-QS effect among cephalosporin antibiotics (CP, CF, and CT) we used OD based microtiter plate assay. First, we quantified MICs of these antibiotics and MIC value showed the tread of CP>CT>CF. This indicates that CP is required in high concentration as compared to CT and CF to kill *C. violaceum*. Further, we tested sub-MICs to find out the concentration that induces a significant anti-QS effect in CV026. The trend for anti-QS activity was as follows: CP>CT>CF. This tread showed that we need a higher concentration of CP to induce anti-QS activity in *C. violaceum* CV026 as compared to CT and CF. The results of the microtiter plate assay indicate that the CF and CT showed anti-QS activity at lower concentrations. The well assay showed that the CP has the highest anti-QS zone against *C. violaceum*. This might be because the MIC is high for CP and the antibiotic is available in high sub-MIC for bacterial cells. CP, CF, and CT showed comparable anti-QS activity in the screening assay, therefore based on our results we excluded carbapenems and selected cephalosporins (CP, CF, and CT) for further anti-virulence activity analysis

### Molecular docking supports the competitive binding of cephalosporins to the CviR binding pocket

Computational biology provides powerful tools to understand ligand-receptor interactions at the atomic level [65]. Molecular docking, in particular, can characterize the molecular interactions between small molecules and proteins [66]. We used this technique to gain insights into cephalosporin-CviR receptor interactions. The active site of CviR is occupied by its natural ligand (C10HSL) and it is buried deep inside the ligand-binding domain of the CviR receptor. The receptor forms hydrogen bonds with Tyr80, Trp84, Asp97, and Ser155 and forms alkyl interactions with Met72 and Tyr88 **(Fig 1J-1M)**. The active site is not fully accessible to cephalosporins due to their relatively large size. However, molecular docking revealed that the cephalosporins might have the affinity to bind with amino acids at the edge of the natural binding pocket including Met72 and Tyr88 **(Fig 1J-1M)**. This binding might block ligand binding or may induce conformational changes to block activation of CviR. Molecular docking results showed that all three antibiotics formed H-bonds with Met72. CT also formed hydrogen bonds with Tyr88 and two more amino acids. However, CF and CT also showed unfavorable interactions with Ser89 and Ala94. The docking score predicted the affinity of cephalosporins against CviR (higher to lower) as CT>CP>CF. Our experimental anti-QS trend followed the sequence CP>CT>CF. In both cases, CF showed the lowest activity, supporting the conclusion that CF binding is the weakest of the three ligands. However, in molecular docking experiments, CT showed the highest score. This indicates that there might be other factors including conformational dynamics and drug bioavailability that may contribute to the anti-QS activity. Nevertheless, molecular docking analysis supports the conclusion that QS inhibition by cephalosporins is related to their ability to bind the QS receptor (CviR). This would likely disrupt the receptor’s interactions with the natural ligand. Competitive inhibition is therefore likely the mechanism of inhibition of QS dependent violacein pigment production in CV026.

### Sub-inhibitory concentration of cephalosporins converts the biphasic growth of PAO1 into a monophasic growth

For all three antibiotics, i.e., CP, CF, and CT, PAO1 growth was inhibited at the MIC and unaffected at zero and MIC/4 concentrations. However, at sub-MIC concentrations of MIC/2, growth became monophasic, deviating from the biphasic growth. The biphasic growth was absent for QS-mutant strains (**Fig 2D**). We fitted the growth curves with the equation [67]:

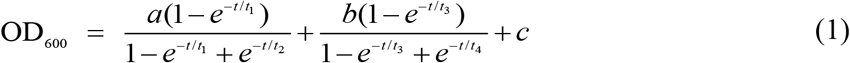

Equation (1) fitted (solid lines) data points (symbols) well (**Fig 2A-2D**). Typically, a monophasic growth described by a logistic equation is observed in a bacteria culture. There are three distinct phases: (1) a lag phase with very slow growth when the bacteria undergo preparatory steps for the next phase, (2) a log phase of exponential growth when the bacteria divide at a constant rate, (3) a stationary phase when the bacteria stop dividing due to unfavorable conditions such as nutrient scarcity. At the stationary phase, *P. aeruginosa* leverages QS pathways and responds to nutrient scarcity by producing virulence factors such as proteases [68] and pyocyanin [69, 70] to trigger a multipronged survival strategy. Proteases can break down large protein molecules available in the extracellular space to smaller components to be used as nutrients [68], whereas pyocyanin can kill competitive bacteria or organisms [71]. As a result, a new phase of growth is facilitated by QS pathways resulting in the observed biphasic growth (**Fig 2A-2C**). This conclusion is supported by the absence of the second growth phase in all QS-mutant strains of PAO1 (**Fig 2D**). An analysis of the measured stationary phases for QS-mutant reveals that the ΔLasR mutant has a stationary phase with the smallest slope, whereas the ΔPqsA mutant has a stationary phase with the largest slope. **Fig 2D** suggests that the inhibition of QS by the four mutants follows the order: ΔLasR>ΔPqsR>ΔRhIR>ΔPqsA. This order agrees with the known hierarchical order of QS pathways in *P. aeruginosa* [72]. The first phase of the growth curve also revealed an interesting observation. The QS-mutations did not affect the first phase of growth and matched well with the first phase of PAO1, suggesting that the first phase is not controlled by QS pathways. For cephalosporins, CP and CT showed the least and most effects respectively on the preparatory lag phase of the first monophasic growth. The MICs for CP, CF, and CT against PAO1 were measured to be 2 μg/mL, 4 μg/mL, 32 μg/mL respectively, i.e., CP is the most potent among the three against PAO1 (**Supplementary Fig 6**). Despite the low MIC, sub-MIC concentrations did not delay the start of the exponential log phase of PAO1 growth, suggesting that CP induced less environmental stress to PAO1 leading to a shorter preparatory lag phase. 1/2^th^ MIC concentration was significantly affecting the growth of the cells and delaying the log phase, while 1/4^th^ MIC did not affect the growth profile of the bacteria, and cells were effectively growing in presence of these concentrations of antibiotics (**Fig 3A-3F**). Therefore, 1/4^th^ MIC concentration of each antibiotic was selected to test anti-virulence and anti-biofilm activities against PAO1. These findings are in agreement with previous reports that show these antibiotics alter gene expression induced mutations and alter growth profiles at sub-MICs leading to growth inhibition of bacteria [73, 74].

### Pyocyanin production was blocked by sub-inhibitory concentrations of cephalosporins

Pyocyanin is a blue colored secondary metabolite and major virulence factor responsible for the extreme toxicity of *P. aeruginosa* against host cells [75]. Previous research has shown that the pyocyanin deficient phnAB and PhzB1 mutants produce a low level of pyocyanin leading to reduced mortality in the burn wound model of *P. aeruginosa* [76, 77]. Pyocyanin also induces pro-inflammatory and pro-oxidant responses leading to host cell lysis [78]. It attacks neutrophils and epithelial cells by reactive oxygen species (ROS) induced apoptosis [79]. Therapy targeting pyocyanin production in *P. aeruginosa* may suppress the tissue damage during infection. Our results showed that all three antibiotics significantly suppressed the pyocyanin production *in vitro* and their ability in the suppression of pyocyanin production was as follows: CP>CF>CT (high to low) (**Fig 3A and 3B**). CP showed the highest effect as compared to CT and CF. Previously anti-QS effect of macrolide antibiotics (azithromycin and erythromycin) has been related to pyocyanin inhibitory effect [80–82]. The *in vitro* anti QS activity followed the pattern as follows: CP>CF>CT (high to low) and *in vitro* pyocyanin production also followed the same pattern (CP>CF>CT). This indicates that the QS inhibitory potential of cefepime targeted the pyocyanin production in *P. aeruginosa* PAO1.

### Inhibition of QS by cephalosporins is accompanied by reduced motility of PAO1

Cellular motility is responsible for the spread of infection and assists bacterial cells to translocate toward nutrient-rich environments [83]. *P. aeruginosa* is a motile Gram-negative bacterium and motility phenotypes significantly influence the establishment and spread of infections. The effect of cephalosporins on swimming motility showed the following pattern: CP>CF>CT **(Fig 4A_i_ and Fig 5A).** We already mentioned that the QS mutant strains showed no significant difference in the swimming motility suggesting there is no relationship between swimming motility and QS. Interestingly, CP, CF, and CT antibiotics showed motility inhibition of wild type PAO1 and its isogenic QS mutant strain. This provides a clue that cephalosporins might have a direct effect on flagellum proteins. Flagellum proteins propel bacteria and allow swimming motility behavior, which helps in initial attachment, surface coverage, and biofilm dispersal [84]. Compound targeting swimming motility may inhibit surface colonization leading to the defective biofilm formation. Inhibition of swimming motility by cephalosporins may target surface colonization and biofilm dispersal by *P. aeruginosa*. Previous research has shown that the antimotility compounds target gene synthesis of flagellum and pilus synthesis to inhibit the movement of *P. aeruginosa* [85]. Targeting gene expression or functions of flagellum molecular motors might be the key mechanism of swimming motility inhibition.

Swarming motility is also a flagella-dependent movement of cells to spread as a biofilm over biotic and abiotic surfaces [86]. Previous research has shown that QS controls biofilm development by regulating swarming motility phenotypes [87]. Our results also showed that swarming motility is dependent on las and rhl signaling systems and independent of the PQS signaling system. Cephalosporins similar pattern in swarming motility inhibition (CP>CF>CT) (**Fig 4B_i_** and **Fig 5B**). However, there was a significant difference in the swarming motility of ΔLasR and ΔRhlR mutant strains with and without cephalosporins. This indicates that cephalosporins might also have a direct effect on the flagellar motility. Moreover, the inhibition of swarming motility also provides strong evidence that cephalosporins may interfere with the QS communication of *P. aeruginosa*.

Twitching motility is a flagella-independent bacterial motion on moist surfaces by extension and retraction of polar type IV pili of *P. aerugionsa* [88]. Previously, researchers debated the dependence of twitching motility on QS pathways [89, 90], however, it is known that twitching motility is required for the establishment of infection and biofilm formation. Our results showed that las and rhl quorum sensing systems were involved in pilli-mediated twitching motility. The pattern of twitching motility inhibition by cephalosporins was similar to swimming and swarming motility inhibition pattern (CP>CF>CT) (**Fig 4C_i_** and **Fig 5C**). There was also a significant difference in the twitching motility of ΔLasR and ΔRhlR mutant strains with and without cephalosporins suggesting their alternate inhibitory mechanisms. In contast, CT showed no effect on twiching motlity phenotypes of PAO1 or its isogenic mutant. This indiates that the cephalospoinrs except CT has direct effect on pilli function. The future research require indepth understanding of the molecular mechanism of antibiotics on flagellar and pilli proteins and quantifying the effect on cell movements.

### Inhibition of QS by cephalosporins is accompanied by reduced biofilm formation by PAO1

Bacterial biofilms are complex three-dimensional bacterial colonies that are highly resistant to antibiotic treatment [91]. The las QS system is required for the maturation of *P. aeruginosa* biofilms [92] and plays an important role in later biofilm development stages. The rhl QS also plays a key role in biofilm formation and research also has shown that the rhl can control biofilm formation independent of its canonical HSL autoinducer activation pathway [93]. The PqsR mutation has been linked with reduced biofilm formation in *P. aeruginosa* [94]. The regulatory role of las, rhl, and pqs system in biofilm formation provides an exciting opportunity to develop anti-QS therapy as potential antibiofilm agents. Our results clearly showed that the anti-biofilm effect of cephalosporins at sub-inhibitory concentrations. SEM analysis and quantitative live bacterial count showed the following pattern of antibiofilm efficacy of cephalosporins: CP>CF>CT **(Fig 6).** This pattern was similar to our anti-QS activity pattern. In addition, despite the low anti-QS activity, the inability of CT to inhibit twitching motility might also contribute to its low antibiofilm efficacy. CP was the most effective antibiofilm agent and significantly inhibited seven-day-old biofilms. CF was ineffective against day-five and CT was ineffective against day-five and day-seven old biofilms. The biofilm formation was observed in the presence of antibiotics and in the future, it would be interesting to monitor the biofilm eradication effect on established biofilms. The exopolysaccharide matrix plays a key role in protecting against environmental insult and antimicrobial agents [95]. It also helps the cells during initial attachment and regulates the transition from reversible to irreversible attachment biofilm development [96]. Cephalosporin treatment has significantly reduced the exopolysaccharide matrix production in early to late biofilm stages. SEM analysis showed that the CP was found to be the most effective treatment to reduced exopolysaccharide matrix production even in late biofilm stages. In the future, quantitative estimation of alginate, extracellular DNA, and matrix proteins might precisely quantify the comparative effect of cephalosporins on exopolysaccharide matrix production. Our results suggested the pattern of inhibition as follows: CP>CF>CT (high to low). CP showed the highest antibiofilm activity against *P. aeruginosa* biofilms.

### Cefepime enhanced antimicrobial efficacy of aminoglycosides against PAO1

A combination of treatments against bacterial pathogens is preferred due to its high potency. Various studies have shown the synergistic activity of antimicrobial agents against *P. aeruginosa*. Although their mechanism of synergistic activity is poorly understood. Previous studies have shown that the combination of CF with tobramycin increases the antimicrobial potency against *P. aeruginosa* [97]. The synergistic activity of gentamicin and ciprofloxacin has also been reported against *P. aeruginosa* [98]. In addition, CP has shown synergistic activity with amikacin against Gram-positive and Gram-negative pathogens [99, 100]. Our results showed that CP has significantly decreased the MIC of aminoglycosides **(Fig 7)**. There was 75% decrease in kanamycin MIC (from 32 μg/mL to 8 μg/mL) and streptomycin MIC (from 4 μg/mL to 1 μg/mL) in presence of CP. Neomycin, gentamicin, and tobramycin showed a 50% decrease in MIC values in presence of cefepime. Our results suggested that although CP enhanced antimicrobial activity of all the tested aminoglycosides however, it worked better with kanamycin and streptomycin **(Fig 7)**. Further research is required to validate its efficacy in animal models to confirm the *in vitro* results.

### Cephalosporins may bind to *P. aeruginosa* QS receptors to inhibit ligand binding

The docking score of cephalosporins to LasR showed the following pattern: CT>CP>CF. This docking pattern was similar to the docking score of CviR, and CT showed the highest binding affinity to the LasR receptor. The LasR docking scores are low as compared to the CviR receptor docking scores indicating a low binding affinity for LasR. The LasR natural ligand-binding pocket is buried deep inside the receptor and hence most likely inaccessible to the cephalosporins due to their large size **(Fig 8)**. However, the antibiotics showed significant interactions with amino acids surrounding the binding pocket. This suggests that the cephalosporins might be able to block the entry of the ligands to the binding pockets thereby inhibiting the activation of LasR. CT and CF showed single unfavorable interactions with the LasR receptor. CP, CT formed two hydrogen bonds, and CF formed three H-bonds without any unfavorable interactions with the LasR receptor. Interestingly, CT showed the highest docking affinity with the lowest anti-virulence effect *in vitro*. This highlights the fact that in addition to the anti-QS effect cephalosporins possess alternative anti-virulence pathways to suppress the virulence of *P.aeruginosa*.

Our docking results with PqsR showed the following pattern: CP=CT>CF. All three cephalosporin showed high docking scores with the receptor. The docking sores of CP and CT were found to be equal. However, CF showed one unfavorable interaction and that might have contributed to its slightly lower docking score as compared to CP and CT. Moreover, the docking score of QZN was −7.4 and ligand was not forming any hydrogen bonds with the receptor. CP formed a hydrogen bond with leu208 and π-interactions with Ile236 **(Fig 8)**. It is important to note that these amino acids were also contributing towards alkyl interactions with QZN indicating CP was able to bind to the active site of LasR. Similarly, CT formed hydrogen bonds with Ile236, and CF formed alkyl interactions with Ile236, which is also shared by QZN. This clearly indicates that all three antibiotics were interacting with a common target area on PqsR. The high docking score of CP with PqsR was consistent with its potent pyocyanin inhibition activity. Previous reports have shown that the pqs system controls pyocyanin production. Our results indicated that cephalosporins have a high affinity for PqsR. The high predicted binding affinities for cephalosporins with LasR and PqsR indicate that their binding plays an important role in suppressing QS regulated virulence expression.

### Sub-inhibitory concentration of cephalosporin reduced virulence potential of *P. aeruginosa* in *C. elegans*

*C. elegans* has been widely used as an efficient animal model system to investigate anti-infective activities of synthetic and natural compounds against various bacterial pathogens [101, 102]. In addition, *C. elegans* has been used as an established animal model to investigate the virulence of *P. aeruginosa* [103]. *P. aeruginosa* mediated worm mortality is induced by cyanide asphyxiation and paralysis [104]. The cyanide production in *P. aeruginosa* is controlled by the *hcn* operon and regulated by LasR and RhlR QS systems [105]. QS systems have been linked to play a crucial role in *C. elegans* infection. Our results were in line with previous findings that suggest the high virulence of wild type PAO1 strain achieving 100% mortality after 72. On the other hand,we have found that QS mutant strains showed significantly high *C. elegans* surviving score (%) as follows: ΔPqsA (36.7±5.8%) > ΔLasR (31.7±11.2%) > ΔRhlR (27.3±7.6% %) > ΔPqsR (26.7±10.4%). Our results showed that all mutant strains were defective in virulence and *C. elegans* survived the infection by QS mutant strains; however, the protection was not comprehensive since the survivor rate was low with all the mutant strains. ΔPqsA and ΔLasR mutants showed the highest surviving, although the score with ΔRhlR and ΔLasR mutant was also significantly high. Treatment of *C. elegans* with cephalosporins with sub-MICs of cephalosporins has shown a significant protective effect on the worm’s mortality. For CP, CF and CT the worm survival score at 72 hr was 65.3±5.0%, 58.0±4.3% and 48.7±8.1% (CP>CF>CT). Cefepime (CP) showed the highest protective effect in terms of *C. elegans* survival as compared to CF and CT. The high protective effect of CP may be linked with its high anti-QS activity and anti-virulence efficacy against *P. aeruginosa*. Previous research has shown that QS inhibitors protect *C. elegans* and human lung epithelial cells from *P. aeruginosa* [106]. Anti-QS effects of clove oil have shown to reduce las- and rhl-regulated virulence factors such as LasB, protease, and pyocyanin production, motility and exopolysaccharide production, and protects C. elegans against *P. aeruginosa* infection [107]. The anti-QS activity of 2, 5-Piperazinedione has been linked in the proteolytic and elastolytic activities of PAO1 and significant protection of *P. aeruginosa* pre-infected *C. elegans* [108]. The anti-QS activity of antagonistic compound phenylacetic acid (PAA) has been shown to suppress QS-dependent exopolysaccharide, pyocyanin, elastase, and protease production in PAO1 and treatment of PAO1-infected *C. elegans* showed enhanced survival after treatment with PAA [109].

It is important to consider that in our study, none of the antibiotics (at sub-inhibitory concentrations) showed complete protective effect at the sub-MIC concentration (at 72 h) indicating that the anti-QS therapeutics may protect from acute infection progression, however for complete treatment of infection combinatorial therapies are needed. Further research is needed to identify the pathways or strategies to ‘switch off ‘the virulence mechanism leading to complete protection from the infection.

## CONCLUSIONS

In the present study, we have shown that cephalosporins can be used as QS inhibitors. We proved its anti-virulence potential in a *C. elegans* PAO1 infection model. Based on the *in-silico, in-vitro* and *in-vivo* results although all three potent inhibitors showed significant therapeutic effects, however, cefepime was proved the best in terms of virulence suppression, biofilm inhibition and protection of *C. elegans* against *P. aeruginosa* infection. Further, targeting *P. aeruginosa* QS pathways suppress virulence factor production and enhance its susceptibility of *P. aeruginosa* to aminoglycoside antibiotics. In addition, the antibiofilm efficacy of cephalosporins may prove to be a valuable finding to develop novel treatment strategies against biofilm-associated *P. aeruginosa* infections, especially in cystic fibrosis patients.

## MATERIALS AND METHODS

### Bacterial strains used in the study

*Chromobacterium violaceum* CV026, *Pseudomonas aeruginosa* wild type strain PAO1 and its defective QS mutant strains including ΔLasR, ΔRhlR, ΔPqsA, ΔPqsR were kindly provided by Dr. Paul Williams (Professor of Molecular Microbiology, Faculty of Medicine and Health Science, University of Nottingham, United Kingdom). *Chromobacterium violaceum* CV026 was cultured using Luria Broth with kanamycin at 50 μg/mL (30°C) under stationary conditions and used for the detection of acyl-homoserine lactone and screening of anti-QS activity of β-lactam antibiotics. All bacterial strains were maintained as 25% glycerol stocks and stored at −80 °C. Fresh subculture was performed from frozen stocks and used for each experiment.

### Antibiotics and chemicals

The following antibiotics cefepime hydrochloride (#PHR1763-1G:LRAB8503, sigma), ceftriaxone disodium salt hemi-heptahydrate (#C5793-1G: 027M4799, sigma), ceftazidime (#CDS020667-50MG:B02487097), oxacillin sodium salt (Cat#28221-5G:097M4853V), imipenem (#PHR1796-200MG:LRAB7177, sigma), doripenem hydrate (#SML1220-50MG:025M4717V, sigma), meropenem trihydrate (#USP-1392454: 50K434), ertapenem (#CDS022172-25MG:B02467020) were procured in dry powder and stored in 4°C under desiccated conditions. Master stocks of 1024 μg/mL were made in sterile distilled water and stored in −80°C for future use. Every time fresh stock of antibiotics was used for experiments. All the other chemicals are of ACS analytical grade.

### Screening of anti-QS activity of antibiotics

#### (a) Agar well diffusion assay

Agar well diffusion assay using *Chromobacterium violaceum* CV026 as biosensor strain was performed to screen anti-QS activity of antibiotics. 50 μL of C6-HSL stock solution (10 mg/mL) was mixed with 10 mL of an overnight grown culture of *C. violaceum* CV026 (OD_600_=1.0) and designated as AHL-CV026 solution. To prepare media plates, 19 mL of tryptose soya agar (1 %) was slowly mixed with 1 mL of AHL-CV026 solution (avoid air bubble formation). Plate the mixture in a Petri plate and let the media cool down until agar is solidified. Use sterile pipette tip (200 μL) to puncher holes in solid agar. Press the pipette and twist to take the punched agar and invert the plate to remove the agar block. Discard the pipette tip and use fresh tips every time to make clean wells. Load different concentrations of antibiotics in each well and incubate the plates for 15 hr at 37°C in a bacteriological incubator. Stock solution (1024 μg/mL) of antibiotic was 10 fold diluted and 50 μL of each dilution was loaded in each well. The amount of antibiotics in well-1 to well-8 was as follows, well-1= 51.2 μg; well-2=25.6 μg; well-3= 12.8 μg; well-4=6.4 μg; well-5=3.2 μg; well-6= 1.6 μg; well-7=0.8μg and well-8= 0.4 μg. After incubation, observe plate in light back growth. Clear zone without bacterial growth and no pigment indicated antimicrobial zone and hazy zone with bacterial growth but without pigment production indicate anti-QS activity of antibiotics. The picture was taken by Bio-Rad imager and compiled in one image with a scale bar (20 mm). Anti-QS zones was were quantified following equation:

> AQS Zone= (**D** _Growth+ No Pigment_ - **D** _No Growth/No Pigment_) [AQS Zone= Anti-QS zone (cm), **D** _Growth+ No Pigment_=Diameter of growth zone with no pigment production, **D** _No Growth/No Pigment_ = Diameter of no growth zone and no pigment].

#### (b) Microtiter plate assay

Briefly, a round-bottom microtiter plate well was filled with 100 μL of LB media. The overnight culture of CV026 was inoculated in 10 mL of LB media and incubated for 3-4 hr until the OD600 reached 0.2-0.3. After incubation cells were harvested and washed three times with phosphate buffer saline (0.01 M, pH-7.4). The washed cells were reconstituted in PBS and OD was adjusted to 0.1 (OD600). Stock cell suspension (10 μL) was used as inoculum for each well. C6-HSL at the final concentration of 10 μg/mL was used for pigment production in each well. Antibiotics were serially diluted in the well. After adding C6-HSL and CVO20, the plate was incubated at 37°C for 24 h. After incubation, the OD567 was measured using a plate reader. We have tested sub-MICs of CP, CF, and CT using a microtiter plate assay. The minimum inhibitory concentration was determined using NCCLS (2002) guidelines (explained in section 2.4).

### MIC determination and sub-MIC selection for *P. aeruginosa* PAO1 strain

The MIC of antibiotics was determined according to NCCLS (2002) guidelines against the standard *P. aeruginosa* wild type strain, PAO1. Round bottom 96 Microwell plate was used for MIC determination. Briefly, the stock solution was serially diluted (2-fold) in cation adjusted Muller Hinton broth and 2-8 × 10^5^ was used as inoculum in the total reaction of 100 μL per well in cation Muller Hinton broth. After 15 hr, resulting in no visible growth was selected as the MIC. Sterile media with bacteria and sterile media with antibiotics are served as positive controls and negative controls. The final plate was also read at OD600 to quantify the effect of the antibiotic on PAO1. MIC, 1/2^th^ and 1/4^th^ MIC was tested on the growth curve profile of PAO1. Respective concentrations showing no significant effect on the growth profiles were selected for anti-virulence factors and anti-biofilm assays against *P. aeruginosa*.

### Motility Competence Assay

*P. aeruginosa* Standard strain PAO1 and QS mutant strains (ΔLasR, ΔRhlR, ΔPqsA, and ΔPqsR) were assessed for motility competence in the presence of a sub-inhibitory concentration of CT, CF, and CP.

#### (a) Swarming motility competence

Media plates were prepared with nutrient agar (8 g/L) complemented with glucose (5 g/L) in the presence of sub-MICs (1/4^th^ MIC) of CT CF and CP. Plates were point inoculated with a sterile toothpick by an overnight culture of *P. aeruginosa* PAO1 and QS mutant strains (ΔLasR, ΔRhlR, ΔPqsA, and ΔPqsR). Swarming Motility competence was determined by measuring circular turbid zones after incubation at 37°C for 24 h.

#### (b) Swimming motility competence

1% tryptone, 0.5% NaCl, and 0.3% agarose media plates with sub-MIC (1/4^th^ MIC) of CT CF and CP were prepared. Plates were point inoculated with a sterile toothpick from an overnight culture of *P. aeruginosa* PAO1 and QS mutant strains (ΔLasR, ΔRhlR, ΔPqsA, and ΔPqsR). After incubation at 37 °C for 24 h, swimming motility was determined by measuring the radius of the circular pattern of bacterial migration.

#### (c) Twitching motility competence

Luria agar plates containing sub-MIC (1/4^th^ MIC) of CT CF and CP were prepared. Plates were stabbed with sterile toothpick up to the bottom of the plates by an overnight culture of *P. aeruginosa* PAO1 and QS mutant strains (ΔLasR, ΔRhlR, ΔPqsA, and ΔPqsR). The plates were incubated at 37 °C for 24 h, media layer was discarded and the hazy zone of growth at the interface between the agar and polystyrene surface was stained with 1% crystal violet stain for 10 min. Plates were washed three times with 5 mL of phosphate buffer saline (pH-7.4, 0.1 mM). After drying, the violet-colored zone of bacterial growth was measured.

### Biofilm inhibition estimation of antibiotics

#### (a) Scanning electron microscopy

Biofilms were grown on a Foley catheter (urinary catheter) surface as follows. Urinary catheters were cut into 10 mm pieces under sterile conditions. Catheter pieces were incubated in 40 mL of Luria broth in a 50 mL falcon tube with 100 μL of PAO1 standard strain overnight culture. Catheter pieces were incubated in the presence of sub-MIC of CT CF and CP for seven days under the stationary condition at 37°C. Each group has four subsets of the experimental group for days 1, 3, 5, and 7. Each day catheter pieces were analyzed using Phenom pro-scanning electron microscope.

#### (b) Colony forming units (CFU) quantification

Catheter pieces in different groups were processed as follows for quantification of live bacterial load. Each catheter piece was dipped in 1 mL phosphate buffer saline (pH-7.4, 0.1 M) and bacterial cell were scrapped from the inner and outer surface of the catheter using a sterile scalpel blade. Serial dilution of the sample was performed and plated on Luria agar plates and plated were incubated at 37°C for 15 hr.

### Effect of antibiotics on pyocyanin production

Pyocyanin was estimated using the method of Huerta *et al*. [34] with slight modification as follows. Briefly, PAO1 was grown under different concentrations (10% to 100%) of Luria broth, and different concentrations were tested for pyocyanin production. At the different concentrations of antibiotics (sub-MIC) of CT CF and CP, the production of pyocyanin production was estimated and compared with control. Chloroform was added to cell-free culture supernatant at different concentration and OD at 690 nm of the chloroform layer was measured.

### Determining the combinatorial effect of aminoglycoside and cephalosporins

MIC concentrations of aminoglycosides, namely, gentamicin sulfate (Sigma, Cat# G1914), neomycin sulfate (Sigma, Cat# PHR1491), streptomycin sulfate (Sigma, Cat# S6501), tobramycin sulfate (Sigma, Cat# T1783), kanamycin sulfate (Sigma, Cat# 60615) were determined using standard 96 well plate assay as mentioned previous section in detail. To determine the combinatorial effect of CP with an aminoglycoside, CP at a sub-inhibitory concentration (0.5 μg/mL) in each well was added with different dilutions of aminoglycosides. The MIC was determined by visual detection of bacterial growth and optical density measurement of each well. OD600nm values were plotted as a heat map using Graph pad prism 8.0.

### Molecular docking studies

Molecular docking studies were performed using Autodock Vina (Version 1.5.6) following standard protocols. The crystal structures of CviR (PDB ID: 3QP8), LasR (PDB ID: 3IX3), and PqsR (PDB ID: 4JVI) were downloaded from the protein databank, at http://www.pdb.org. Ligands, ions, water molecules, and small molecules were removed, and the file was saved in pdbqt format using autodock tools. Ligand structure files in PDB format were downloaded from www.rcsb.org. Structures that were not available in PDB format were converted from sdf using an online converter (https://cactus.nci.nih.gov/translate/). The binding site was located by highlighting key amino acids on the receptor protein in autodock tools. Then, a grid box was placed to encase the binding site area, while minimizing the total volume. A config.txt file was created to specify the receptor, ligand, and the x, y, and z dimensional coordinates of the docking site. The coordinates for protein receptors were provided as in **Supplementary Table.1**). For extensive simulation, the ‘exhaustiveness’ option was set as 10.0. Autodock Vina was run and the best docking conformations were visualized using Pymol (Zalman 3D) software and BIOVIA Discovery studio. Docking conformations of each ligand were analyzed by examining their total energy score. Docking scores of cephalosporin antibiotics were compared with the natural ligand of each receptor.

### Nematode Slow Killing assay

Nematode infection model [110, 111] was used to access the anti-virulence effect of cephalosporins. *C. elegans* N2 strain was procured for CGC. The worms were cultivated in an NGM Petri plate with a bacterial lawn of OP50 cells. L2 and L3 stage nematode (20-40 worms) were transferred to 10 NGM plates to grow enough worms for the experiment. After the 4-5 days growth, the worms were collected in liquid NGM media and the concentration of worms was adjusted to approx. 100 worms/mL. *P. aeruginosa* PAO1 was grown at 40% LB media at 37°C stationary conditions with and without sub-inhibitory concentrations of CP, CF, and CT. Briefly, the young adult nematodes were infected with PAO1 (control and antibiotic-treated) and incubated at 25°C for 15 h. After the incubation, worms were centrifuged to remove free bacteria and transferred to fresh liquid NGM media with and without the sub-inhibitory concentration of cephalosporins. QS mutant strain was also collected and processed similarly to evaluate its effect on nematode killing. Worms were confirmed dead if they did not show any moment under the microscope. Every 15 h, worms were immediately observed under a light microscope for the mortality estimation. The number of dead worms was counted and percentage mortality was calculated as follows: Survival (%) = Total no of worms-Number of dead worms/Total no of worms X100.

### Statistical analysis

All experiments were performed in triplicate and repeated on different days. The effect of antibiotic treatment on pigment production, pyocyanin, motility, live cell count in biofilms, and percentage survival/mortality of *C.elegans* in different groups was evaluated using a two-way ANOVA test. p values were calculated and p<0.05 was considered significant. Results were analyzed using Graph Prism 8.0 software. Values were expressed as mean + standard deviation (SD).

**Table 1.** Comparative analysis of molecular interaction of natural ligands and antibiotics with receptor proteins. Amino acids found only in the natural binding pocket are highlighted in red, while amino acids found in the natural and the antibiotic binding pockets are highlighted in green. Amino acids only found in inhibitor binding pockets are shown in black.

| S. No. | Receptor | Ligand | H Bonds | π-π interactions | Alkyl interactions | Unfavorable interactions | Docking score kcal mol<sup>-1</sup>. |
| --- | --- | --- | --- | --- | --- | --- | --- |
| 1. | CviR | C10HSL | Y80, W84, D97, S155 | -- | M72, Y88 | -- | -6.9 |
|  |  | Cefepime | I69, M72 | Q70 | R71, V75, A94 | S89 | -6.3 |
|  |  | Ceftazidime | M72, S89 | -- | L85, A94, M100 |  | -6.1 |
|  |  | Ceftriaxone | I69, M72, Y88, N92 | -- | -- | A94 | -6.5 |
| 2. | LasR | C12HSL | D73, T75, | -- | L40, A50, Y56 | -- | -8.0 |
|  |  | Cefepime | A50, D65 | F167 | I52 | -- | -5.6 |
|  |  | Ceftazidime | G54, N55, Y56 |  | K16, A58 | E62 | -5.9 |
|  |  | Ceftriaxone | K16, N49 | F167 | I52, A58, R61 | D65 | -6.6 |
| 3. | PqsR | QZN | -- | -- | A102, I149, A168, V170, L189, L208, V211, W234, I236, P238 | -- | -7.4 |
|  |  | Cefepime | L208, R209 | I236 | Y258, I263 | -- | -6.7 |
|  |  | Ceftazidime | P210 | -- | I149, A168, L207, V211, I236, Y258 | E259 | -6.2 |
|  |  | Ceftriaxone | E151, I97, L207, I236, Y258 | T265 | L208, I263, K266 | -- | -6.7 |

## ACKNOWLEDGMENT

*P. aeruginosa* PAO1, its isogenic mutant strains (ΔLasR, ΔRhlR ΔPqsR, and ΔPqsA) and *Chromobacterium violaceum* CV026 were kindly provided by Paul Williams, Professor of Molecular Microbiology, and Faculty of Medicine & Health Sciences. This project was funded by the Professional Development Fund (JKS), Slater Foundation Fund (SKS), and COEDIT grant (SKS) at the Colorado School of Mines.

## AUTHOR CONTRIBUTIONS

L.K. initiated the project and designed the experiments; L.K., J.B., and N.B. performed experiments; L.K. and N.B. performed molecular docking, L.K., performed molecular dynamics simulation experiments; L.K., JKS, and S.K.S performed the analysis of MD simulation data; L.K. and S.K.S. designed experiments on synergistic effects of antibiotics; S.K.S. developed quantitative analysis of growth curves; L.K., N.B., J.B., J.K.S, and S.K.S. analyzed manuscript data; L.K., N.B., J.K.S., and S.K.S. wrote the manuscript. All authors edited the manuscript.

## COMPETING FINANCIAL INTERESTS

The authors declare no competing financial interests.

## Supplementary information

**Supplementary figure-1:**
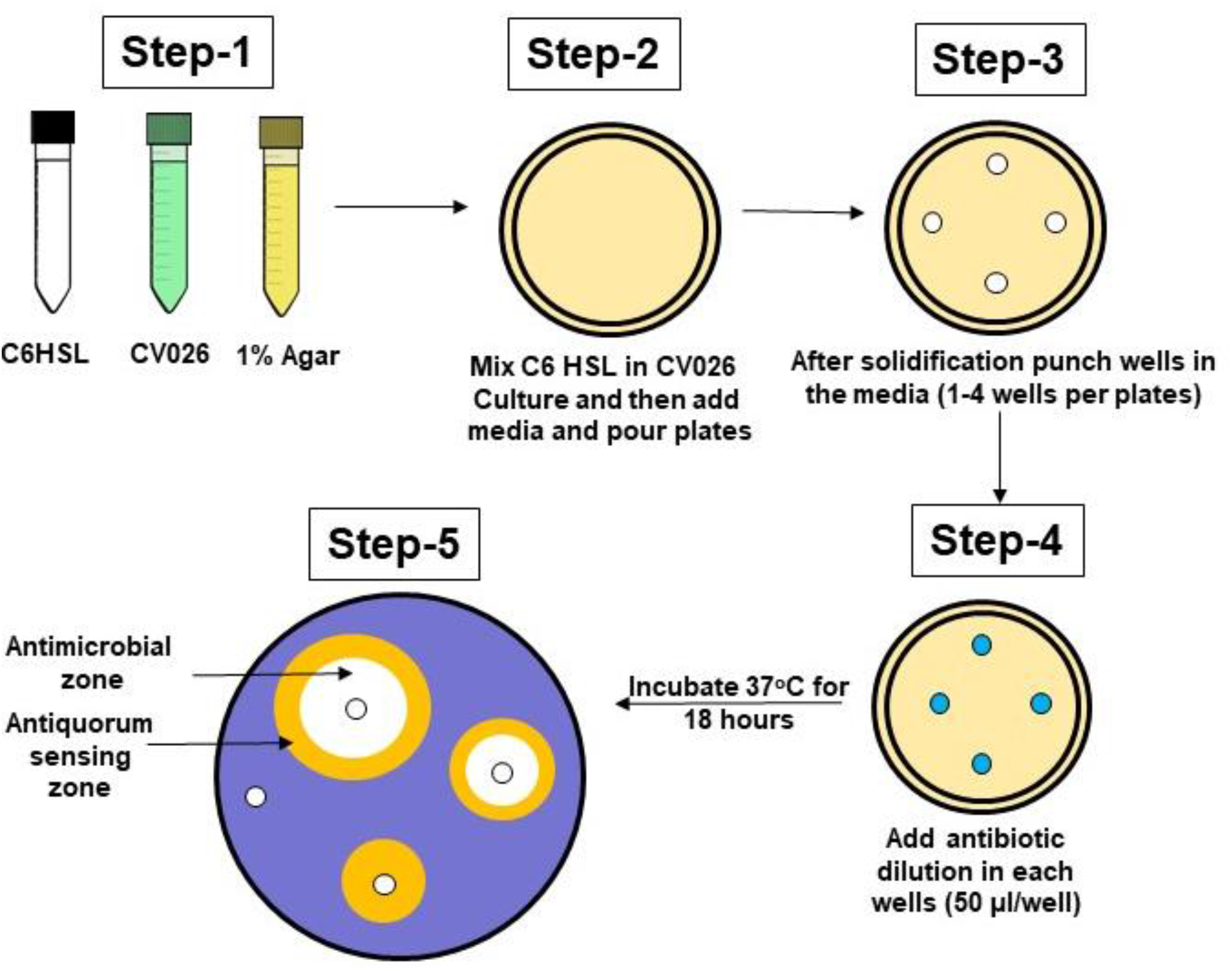
Schematic representation of agar well diffusion assay for the screening anti-QS activity of antibiotics using *Chromobacterium violaceum* CV026 as detector strain. Step-1: Preparation of signal molecule (C6-HSL), overnight *C. violaceum* CV026 culture, and 1% molten agar. **Step-2:** Making agar plates by mixing molten agar, *C. violaceum* CV026, and C6 HSL. **Step-3:** Punching wells using sterile pipette tips. **Step-4:** Filling wells with antibiotic dilutions. **Step-5:** Schematic representation of two distinct zones; antimicrobial zone and anti-QS zones (Interpretation of results of the assay: white zone indicates the antimicrobial zone and yellow zone with bacterial growth with no pigment production indicates anti-QS activity zone).

**Supplementary figure-2:**
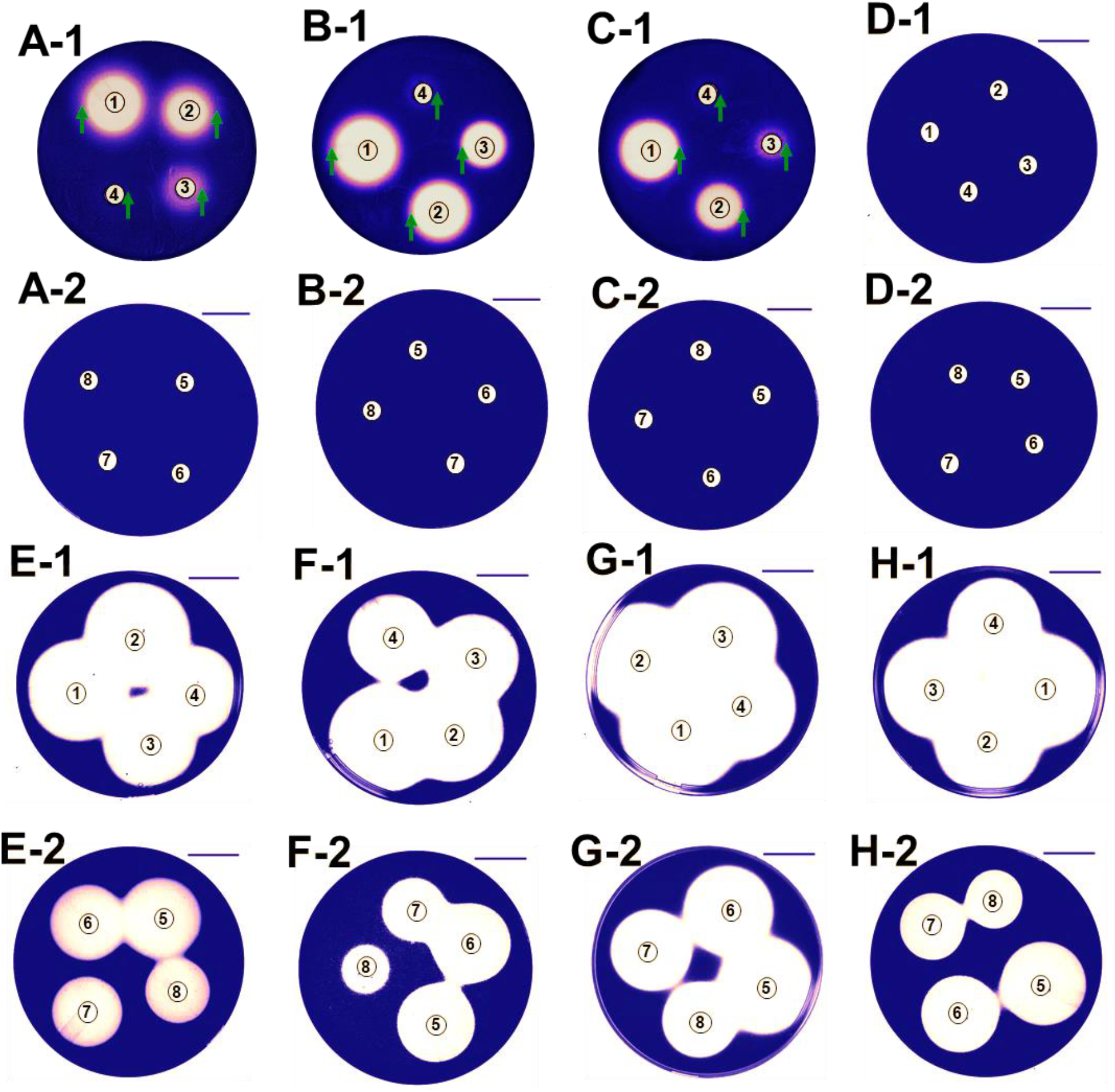
Screening of anti-QS activity of cephalosporin antibiotics using *Chromobacterium violaceum* CV026 as detector strain. Photographs of plates showing zone of growth inhibition and pigment production inhibition (on the edge of the zones) by different classes of antibiotics against *Chromobacterium violaceum* CV026 (Cefepime-A-1, A-2; Ceftazidime-B-1, B-2; Ceftriaxone-C-1, C-2; Oxacillin-D-1, D-2; Imipenem-E-1, E-2; Doripenem-F-1, F-2; Meropenem-G-1, G-2; Ertapenem-H-1, H-2. The green arrow indicates the presence of an anti-QS zone of inhibition (no growth inhibition) for cefepime, ceftazidime, and ceftriaxone (Scale bar 20 mm). In each plate there are eight wells; each well represents different concentration of antibiotics as follows, well-1= 51.2 μg; well-2=25.6 μg; well-3= 12.8 μg; well-4=6.4 μg; well-5=3.2 μg; well-6= 1.6 μg; well-7=0.8μg and well-8= 0.4 μg.

**Supplementary figure 3:**
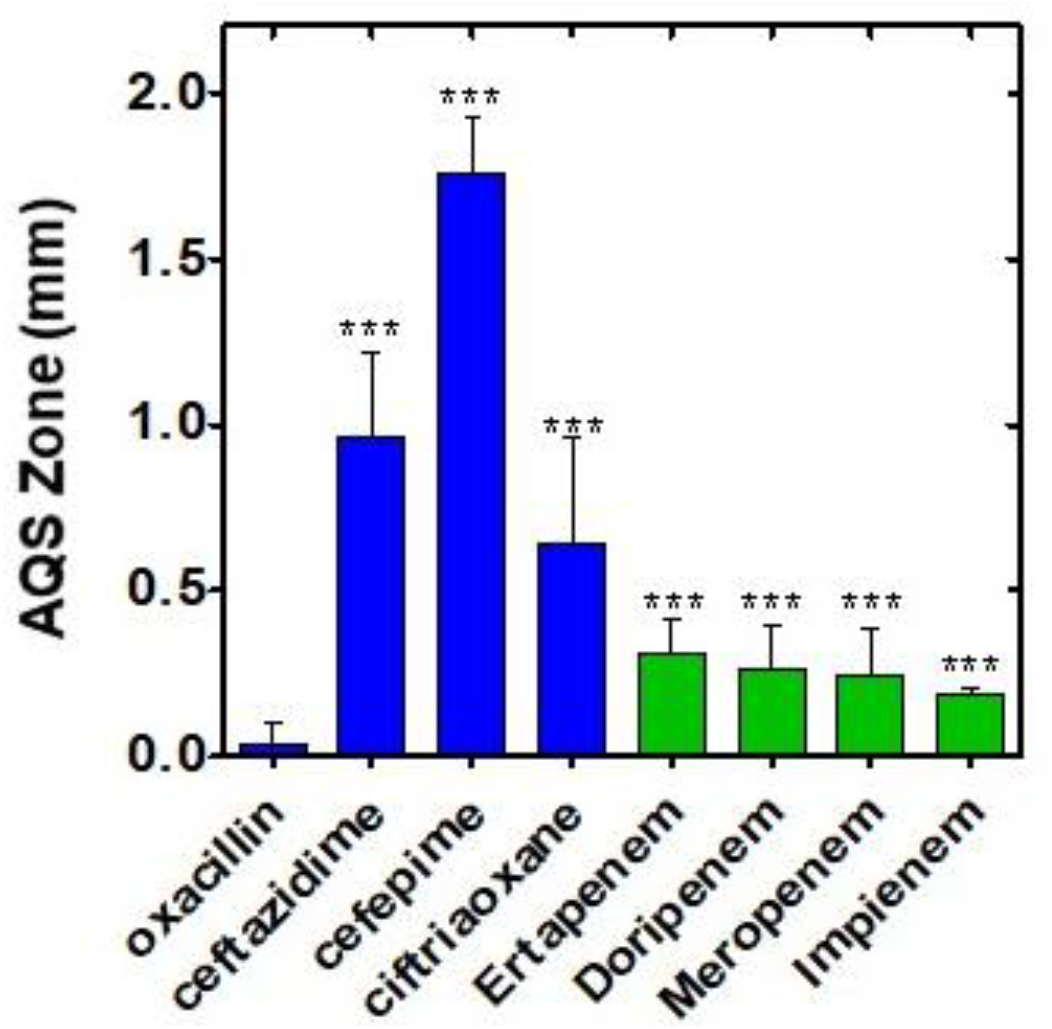
Comparative quantitative analysis of diameters of anti-QS activity of various antibiotics against *C. violaceum* CV026 in agar well diffusion assay. Graphical representation of the anti-QS zone in cm (the area where pigment production was inhibited but the growth of bacteria was present) in agar well diffusion assay. The diameters were compared with the anti-QS zone of oxacillin for the determination of statistical significance. Oxacillin showed no antimicrobial activity against *C. violaceum* CV026. *p<0.05, **p<0.01 and ***p<0.001)

**Supplementary figure-4:**
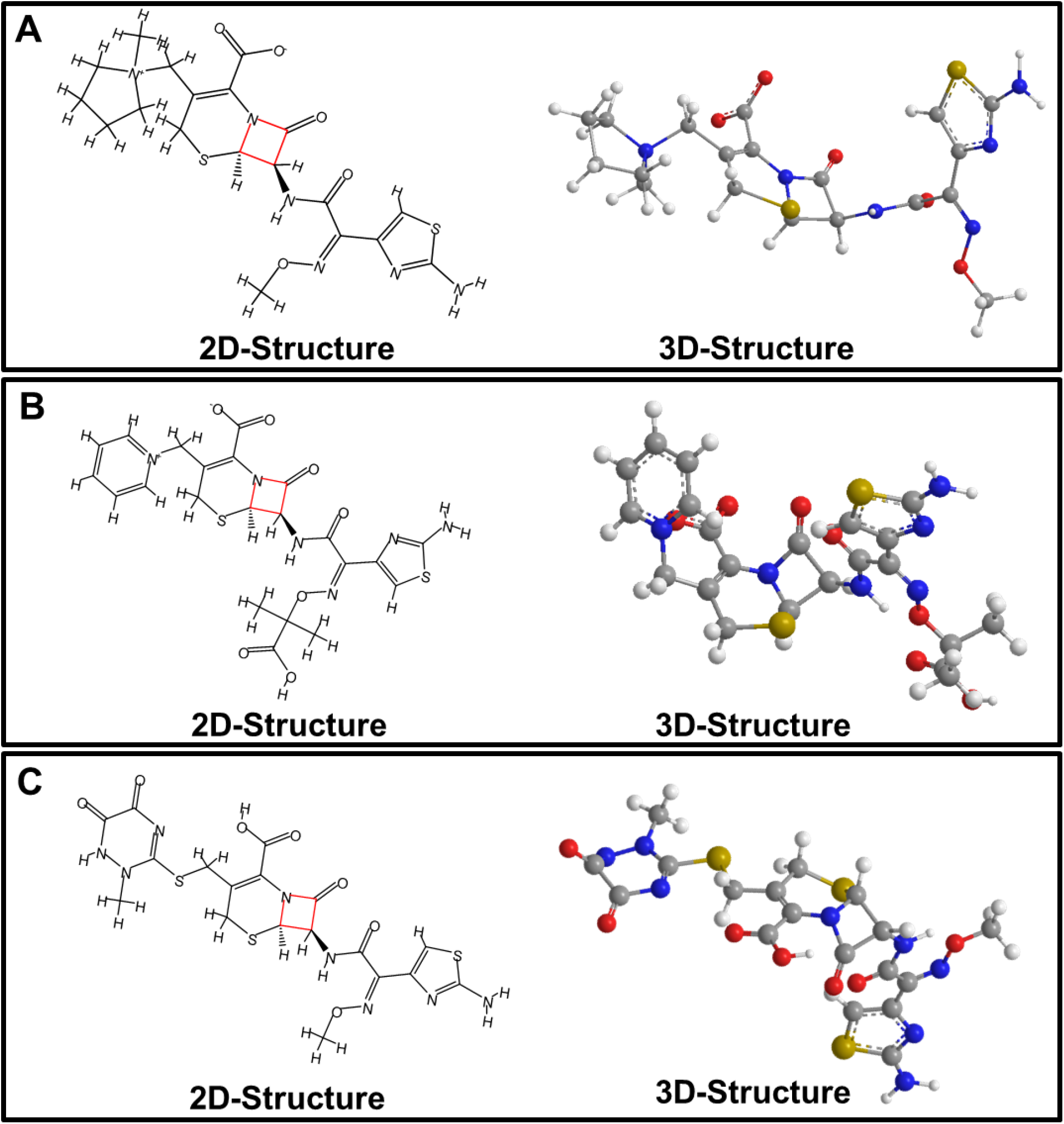
2-D and 3-D chemical structure of cephalosporins; cefepime (A); ceftazidime (B); and ceftriaxone (C) (red square represents the beta-lactam ring of cephalosporin antibiotics. In the 3D structure of antibiotics nitrogen, hydrogen, carbon, oxygen, and sulfur are represented with blue, white, grey red, and light brown balls.

**Supplementary figure-5:**
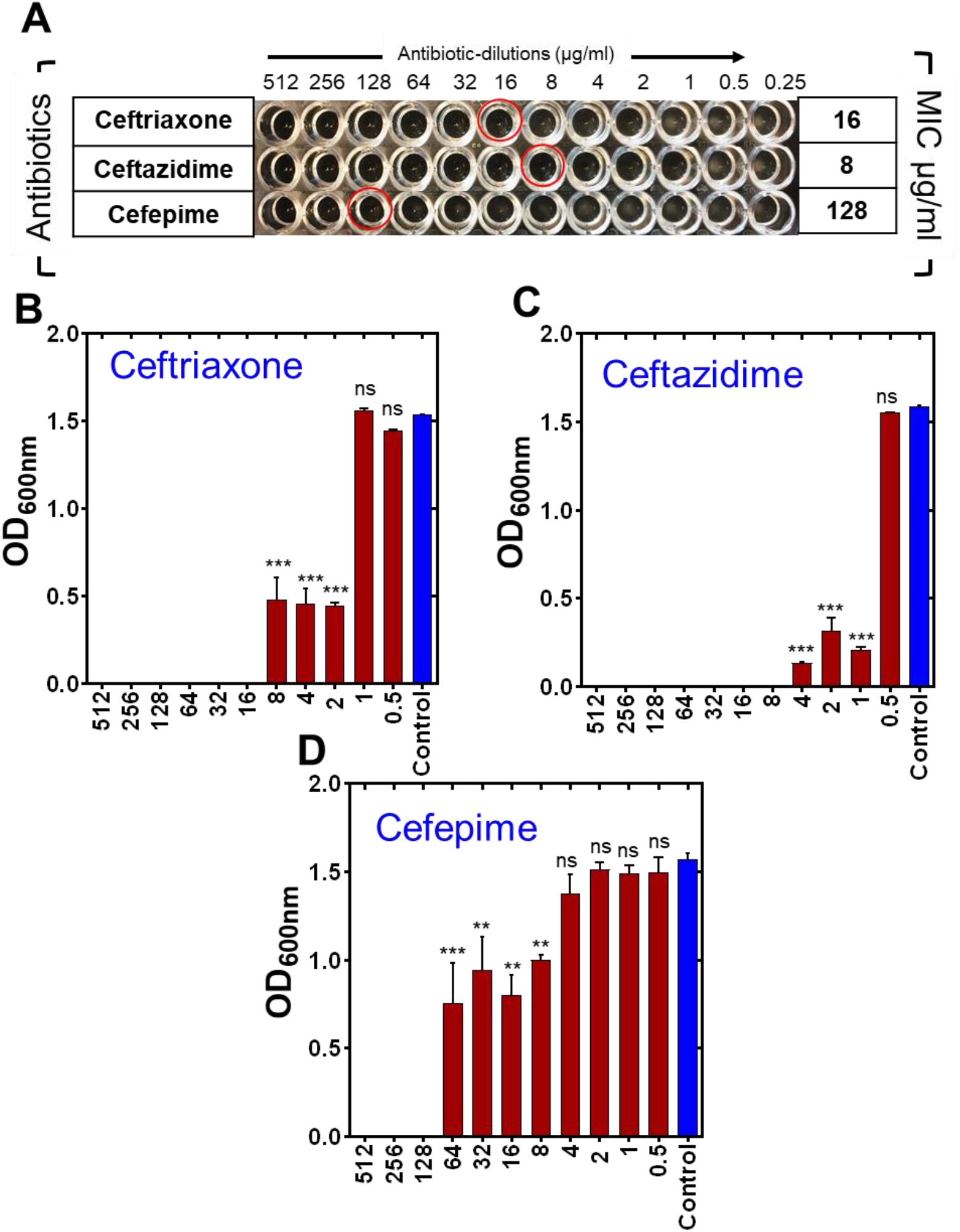
Minimum inhibitory concentration (MIC) determination of cephalosporins against *C. violaceum* CV026. Image of microtiter plate showing wells of PAO1 growth inhibition (A); red circle represents the well of visible growth inhibition. Graphical representation of OD _600nm_ of *C. violaceum* CV026 with various concentrations of ceftriaxone (B), ceftazidime (C), and cefepime (D).

**Supplementary figure-6:**
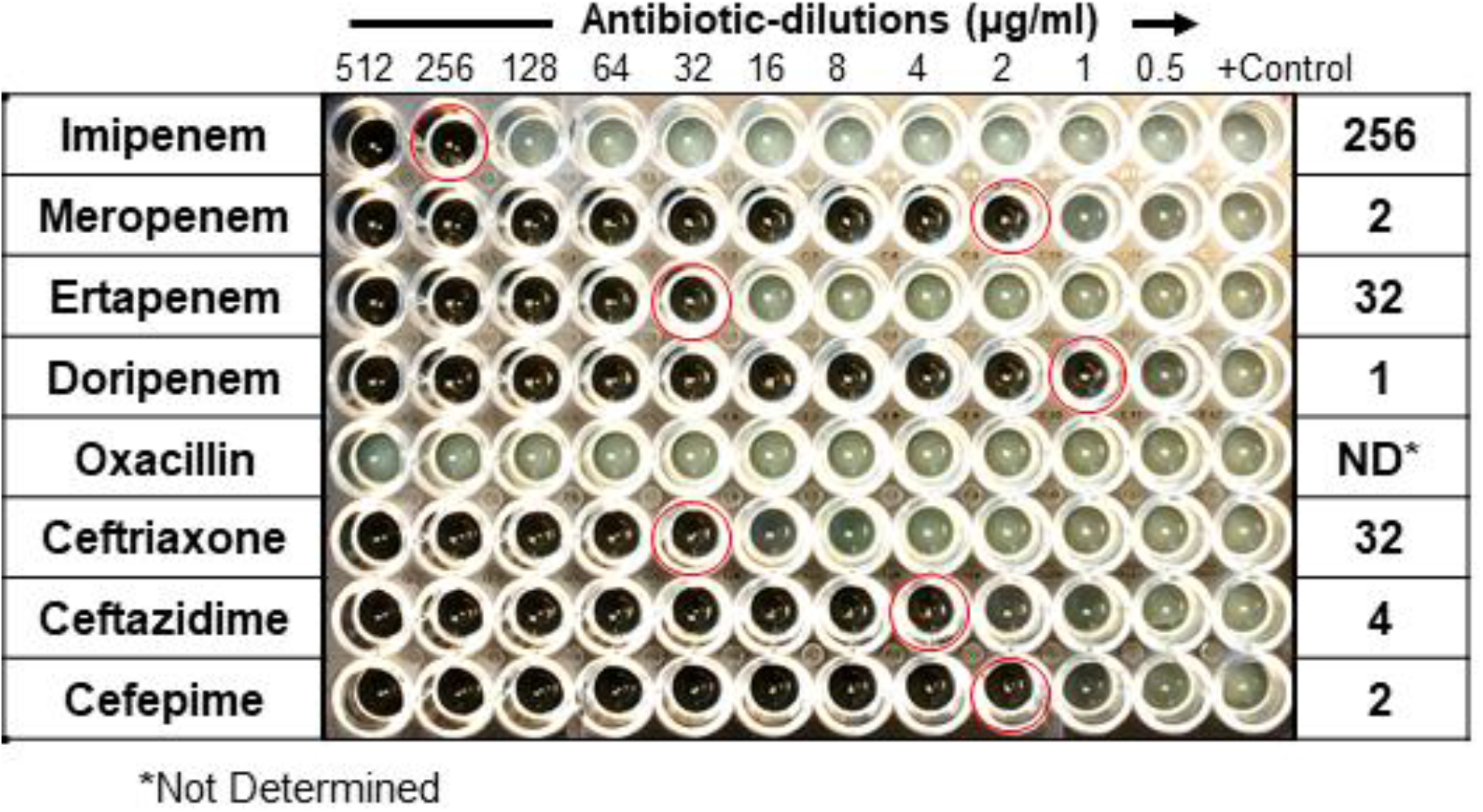
Minimum inhibitory concentration (MIC) determination of *Pseudomonas aeruginosa* PAO1 against cephalosporin antibiotics. Image of microtiter plate showing the growth inhibition. The names of the antibiotics are written on the left panel and the MIC values are written on the right panel of the image.

**Supplementary figure-7:**
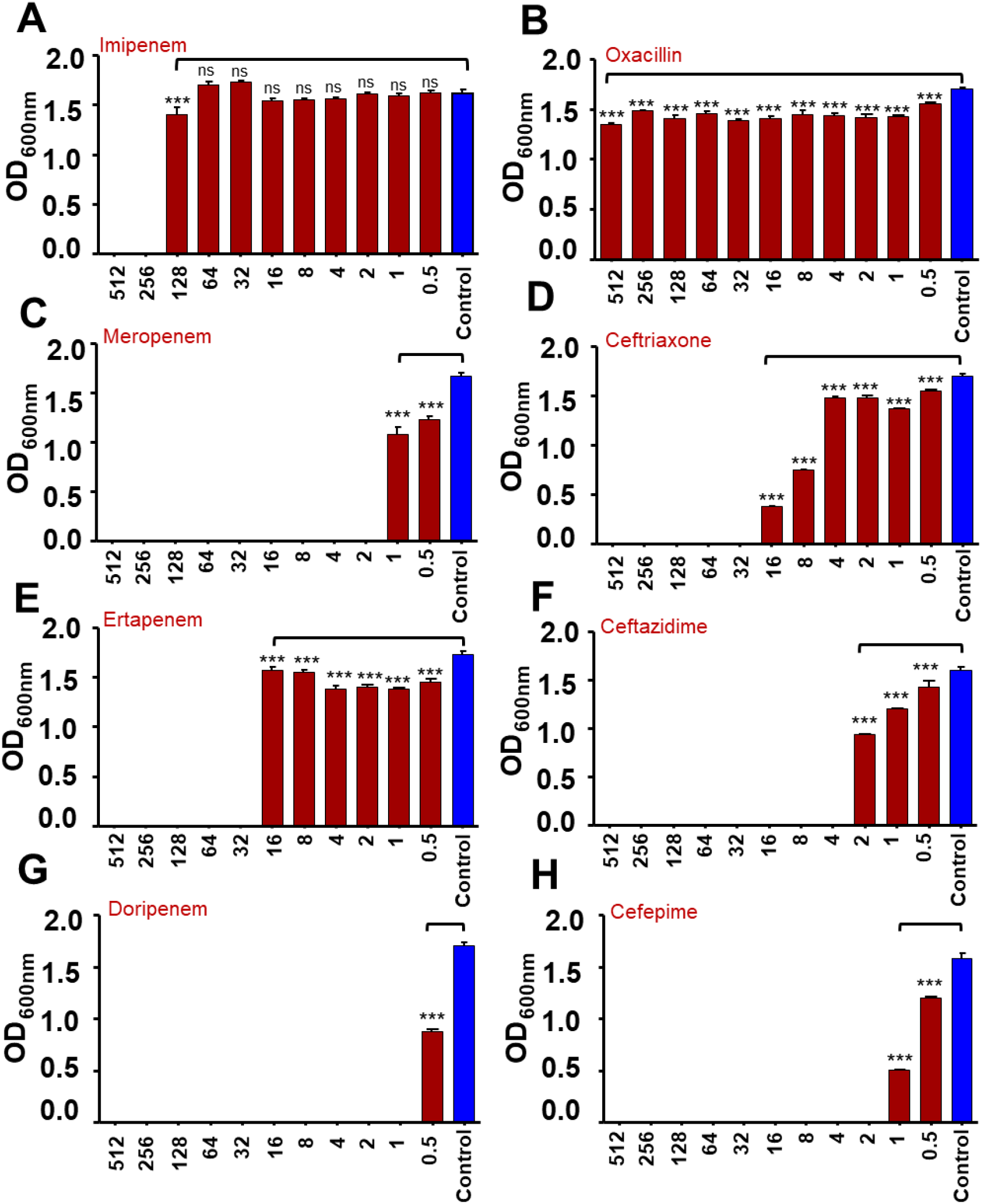
Minimum inhibitory concentration (MIC) determination of *Pseudomonas aeruginosa* PAO1 against Cephalosporins. Graphical representation of OD_600nm_ of *P. aeruginosa* PAO1 at different concentrations of cephalosporins (A-imipenem; B-oxacillin; C-Meropenem; D-ceftriaxone; E-ertapenem; F-ceftazidime; G-doripenem; and H-cefepime).

**Supplementary figure-8:**
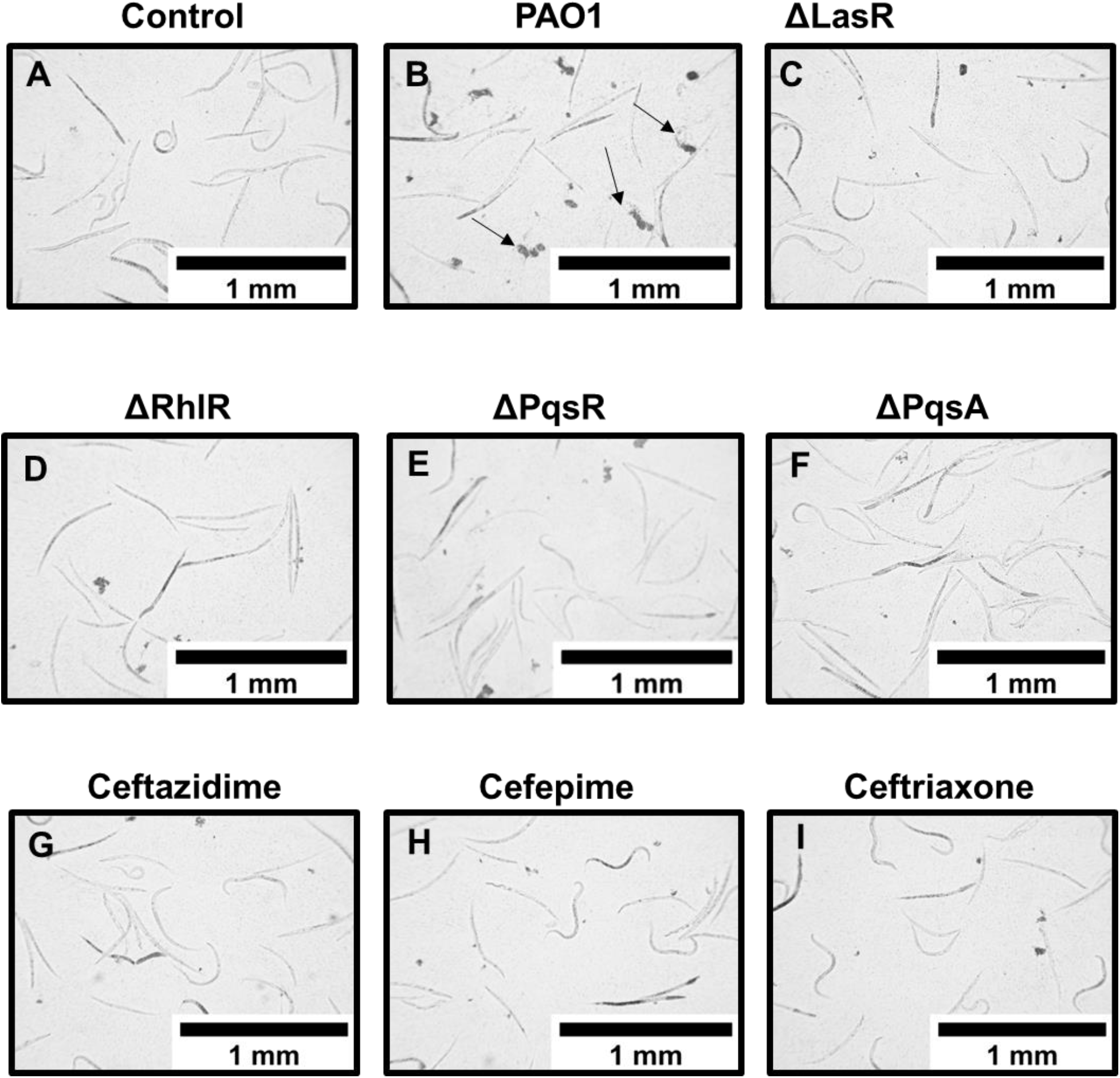
Antivirulence effect of cephalosporins on *C.elegans* survival. Image showing the appearance of *C. elegans* without any treatment (A); after exposing to the supernatant of PAO1 (B); after exposing to the supernatant of QS mutant strains; ΔLasR (C); ΔRhlR (D); ΔPqsR (E); ΔPqsA (F); and exposing to the supernatant of PAO1 grown in presence of a sub-MICs of ceftazidime (G); cefepime (H), and ceftriaxone (I).

**Table 1.** The grid-box coordinates of respective quorum sensing receptors used for molecular docking experiments

| S. No. | Protein | Grid-box coordinates | Value |
| --- | --- | --- | --- |
| 1. | CviR | Center_x | 30.862 |
|  |  | Center_y | 35.116 |
|  |  | Center_z | 18.001 |
|  |  | Size_x | 30 |
|  |  | Size_y | 42 |
|  |  | Size_z | 34 |
| 2. | LasR | Center_x | 9.714 |
|  |  | Center_y | 2.032 |
|  |  | Center_z | 20.29 |
|  |  | Size_x | 36 |
|  |  | Size_y | 30 |
|  |  | Size_z | 28 |
| 3. | PqsR | Center_x | -52.182 |
|  |  | Center_y | 3.466 |
|  |  | Center_z | 10.195 |
|  |  | Size_x | 22 |
|  |  | Size_y | 22 |
|  |  | Size_z | 22 |

**Note:** Exhaustiveness was set to 24 in each simulation.

